# Thalamic Nuclei Differentially Coordinate Propagation of Cortical Slow Oscillations

**DOI:** 10.64898/2026.06.05.730542

**Authors:** Mahmoud Alipour, Wim van Drongelen, Joshua Jacobs, Joel Voss, David Satzer

## Abstract

Sleep slow oscillations (SOs) vary in the spatial extent of their cortical propagation, ranging from widespread Global events to spatially restricted Frontal events. These distinctions could have functional consequences for memory consolidation. It remains unknown whether individual thalamic nuclei are differentially engaged across SO propagation types, and whether thalamic activity before SO onset predicts subsequent propagation. We analyzed simultaneous scalp EEG and thalamic stereo-EEG from 24 full-night recordings in 6 epilepsy patients, sampling 11 thalamic nuclei and enabling direct characterization of nucleus-specific thalamocortical interactions during naturally occurring sleep slow oscillations. Cortical SOs were classified by propagation type and thalamocortical coupling was characterized via peri-event histograms, phase-locking analysis, waveform morphology comparisons, and pre-onset spectral and cross-frequency coupling features. All well-sampled thalamic nuclei showed broad temporal co-occurrence with prefrontal SOs, yet occupied distinct phase positions within the cortical SO cycle. Medial pulvinar (PuM) showed the strongest phase locking across subjects, with significant preference for the ascending post-trough phase. Among the examined nuclei, Frontal SOs were associated with significant phase locking in PuM, whereas Global SOs preferentially synchronized the centromedian nucleus near the cortical down-state trough. Thalamic waveform morphology differed systematically with propagation extent, with opposing effects between pulvinar and intralaminar nuclei. Pre-onset PuM activity showed suppressed alpha, sigma, and beta power and delta-phase cross-frequency coupling before Global SOs (all p < 0.05). The human thalamus engages NREM sleep SOs through nucleus-specific, propagation-sensitive dynamics rather than as a functionally uniform structure. Pre-onset PuM activity contains predictive signatures of cortical SO propagation extent. This finding has implications for closed-loop neuromodulation targeting specific SO states, although replication in larger cohorts is needed.

## 1. Introduction

### 1.1. Cortical Slow Oscillations: From Canonical Synchrony to Propagation Heterogeneity

Slow oscillations (SOs; ∼0.5–1.5 Hz) are a hallmark rhythm of non-rapid eye movement (NREM) sleep, characterized by large-scale synchronized alternations between cortical up-states (periods of intense neuronal firing) and down-states (periods of cellular silence)^1^. These large-amplitude travelling rhythms coordinate neuronal activity across distributed cortical networks and are strongly implicated in sleep-dependent memory consolidation, and synaptic plasticity^2,3^. The prevailing opinion is that SOs emerge intrinsically within neocortical circuits^4^, preferentially involving prefrontal cortex and propagating across the cortex as travelling waves^5^. Within this framework, cortical SO up-states provide temporal windows during which thalamic sleep spindles and hippocampal sharp-wave ripples become coordinated, supporting hippocampal–neocortical memory transfer during sleep^6,7^.

However, this canonical view of SOs as uniform synchronization events obscures a critical layer of heterogeneity. SOs vary substantially in their cortical propagation extent, and these differences carry meaningful functional consequences. Based on scalp EEG spatiotemporal patterns, SOs can be categorized into Global events, which propagate across widespread cortical regions, Frontal events, restricted largely to anterior cortex, and Local events, which occur sparsely without a clear propagation pattern^8^. Crucially, Global SOs exhibit stronger coupling with sleep spindles and are preferentially associated with episodic memory consolidation, whereas Frontal SOs show weaker spindle coordination and no evidence of memory benefit^8,9^. Propagation heterogeneity also carries pathological relevance: individuals with hypersomnolence show an increased proportion of Frontal SOs relative to healthy controls^10^ (Alipour et al., 2025), and intracranial recordings in epilepsy demonstrate that widespread slow waves are preferentially associated with interictal epileptiform activity^11^.

These findings collectively reframe the study of SOs: rather than treating SO occurrence as the primary variable of interest, the spatial propagation structure of individual events emerges as a functionally and clinically significant dimension. What remains poorly understood, however, is how subcortical circuits — particularly those involving different thalamic nuclei — participate differently across SO propagation types, and whether thalamic network states before cortical SO onset already encode information about subsequent propagation. Addressing these questions has direct implications for understanding thalamocortical sleep organization and for the design of predictive closed-loop neuromodulation strategies.

### 1.2. Thalamocortical Coordination of Sleep Slow Oscillations: Nucleus-Dependent Heterogeneity

Although SOs have traditionally been viewed as predominantly neocortical phenomena, converging evidence indicates that the thalamus actively participates in their generation and coordination^12^. Thalamocortical neurons can intrinsically generate slow rhythmic activity through interactions between T-type calcium currents and hyperpolarization-activated cation currents^13,14^, and the thalamic reticular nucleus (TRN) similarly exhibits intrinsic slow oscillatory dynamics^15^. These observations contributed to network-level models proposing that SOs emerge through dynamic interactions across neocortex, thalamocortical neurons, and TRN circuits rather than from a purely cortical generator^16^. Consistent with this view, optogenetic burst activation of centromedian thalamic neurons in mice initiates cortical up-states ^17^, and midline thalamic neuronal bursting activity contributes to synchronized cortical down-state transitions and terminates up-states^18^.

Critically, thalamic involvement in SO dynamics is not uniform across nuclei. Animal studies have demonstrated nucleus-dependent differences in SO phase preference and timing relative to cortical activity^19,20^, and human intracranial recordings have confirmed this heterogeneity: anterior thalamic nucleus SOs precede neocortical SOs, whereas mediodorsal thalamic SOs follow cortical activity^12^. However, human evidence remains limited to a small subset of nuclei, and it is unknown whether thalamic coupling profiles differ according to the spatial propagation extent of the associated cortical SO — a question relevant to understanding whether thalamocortical circuits contribute differently to Global versus Frontal events.

### 1.3. Pre-Onset Network States and the Predictability of SO Propagation

A related unresolved question is whether the thalamic network state preceding SO onset contains information about the subsequent spatial propagation of the cortical event. Closed-loop neuromodulation systems targeting sleep oscillations currently operate primarily in a reactive manner, detecting SOs after onset and thereby limiting temporal precision^21^. Recent scalp EEG work by our team demonstrated that pre-onset cortical activity predicts whether an upcoming SO will propagate globally or remain frontally restricted (Alipour et al., bioRxiv)^22^, suggesting that large-scale network state before SO onset carries predictive information about subsequent SO dynamics. Whether analogous predictive signatures exist within the human thalamus — and whether thalamic recordings could enhance the precision of such predictions — remains unknown. Addressing these questions is important for understanding how thalamocortical networks coordinate sleep oscillations across distributed brain regions and may help inform future closed-loop stimulation strategies targeting specific SO states.

### 1.4. Overview of the Present Study

We investigated thalamocortical SO coupling across multiple thalamic nuclei during human NREM sleep using simultaneous scalp EEG and thalamic stereo-EEG (SEEG) recordings from 24 full-night sessions in 6 patients undergoing presurgical epilepsy monitoring, providing coverage across 11 thalamic nuclei. We first characterized the temporal and phase relationships between thalamic and prefrontal cortical SOs across N2, N3, and NREM sleep (regardless of stage) to determine whether distinct nuclei exhibit different coupling profiles relative to the cortical SO cycle. We then examined whether thalamic SO organization — including timing, phase, and waveform morphology — differs systematically according to cortical SO propagation extent, comparing Global and Frontal events as well as propagation patterns along an anteroposterior gradient. Finally, we tested whether pre-onset thalamic activity contains spectral and cross-frequency coupling signatures predictive of subsequent cortical SO propagation type. Together, these analyses provide a multi-nucleus characterization of thalamocortical SO dynamics in humans and directly link thalamic network states to the spatial propagation structure of cortical slow oscillations.

## 2 Materials and Methods

### 2.1. Participants and Recordings

This study included six adults with drug-resistant focal epilepsy (5 males; mean age 35 ± 9.5 years) who underwent simultaneous scalp EEG and stereo-EEG (SEEG) monitoring. Of 31 available full-night recordings, 24 sessions met the inclusion criteria and were retained for analysis. Sessions were excluded because of substantial recording artefacts, absence of scalp EEG data, or temporal proximity to seizure events; recordings occurring within 5 minutes before or 30 minutes after a seizure were excluded. Written informed consent was obtained from all participants, and all procedures were approved by the University of Chicago Institutional Review Board (protocol 24-1692).

Data were acquired at sampling rates ranging from 1024 to 2048 Hz and subsequently resampled to 256 Hz using the MATLAB resample function. Scalp EEG recordings were obtained using a standard clinical 10–20 electrode configuration comprising 20–26 electrodes per participant and were originally referenced to FCz. For scalp SO detection, signals were transformed to a common average reference (CAR), calculated as the average of all scalp electrodes after excluding mastoid channels, to reduce spatial bias. SEEG recordings were re-referenced using a Laplacian approach in which each contact was referenced to the mean signal of its two adjacent contacts along the same electrode shaft; contacts located at shaft termini were referenced bipolarly. This referencing strategy reduces the influence of volume conduction while maintaining anatomical specificity^23^. To remove line-noise contamination, 60 Hz and 120 Hz notch filters (4th-order zero-phase Butterworth band-stop filters implemented with MATLAB filtfilt) were applied before all subsequent analyses.

### 2.2. Thalamic Electrode Localization

Thalamic SEEG contacts were localized to individual nuclei through co-registration of post-implantation CT scans with pre-implantation MRI, followed by normalization to Montreal Neurological Institute (MNI) space using Brainstorm. Anatomical labels were then determined using the Automated Anatomical Labeling atlas version 3 (AAL3). Based on this procedure, contacts were classified into 11 thalamic nuclei: anterior (ANT), central lateral (CL), centromedian (CM), mediodorsal (MD), parafascicular (Pf), anterior pulvinar (PuA), lateral pulvinar (PuL), medial pulvinar (PuM), ventral anterior (VA), ventrolateral (VL), and ventroposterolateral (VPL). For analyses conducted at the whole-thalamus level, contacts from all nuclei were additionally combined into a single pooled thalamic group. The number of valid recording contacts (contact-session pairs) differed across nuclei. The largest sampling was obtained from PuM (N = 78 contacts from 5 patients), followed by VPL (N = 32), PuL (N = 21), CM (N = 16), VL (N = 8), MD (N = 6), CL (N = 6), Pf (N = 4), PuA (N = 4), ANT (N = 1), and VA (N = 1). Nuclei with only one contact (N = 1) could not support group-level inference and are reported for descriptive completeness only. Recording coverage differed across patients: not all nuclei were sampled in every patient, and the number of recording nights and contacts varied by nucleus (Supplementary Table S1).

### 2.3. Sleep Staging

Sleep staging was performed manually in 30-second epochs by a certified clinical sleep specialist according to American Academy of Sleep Medicine (AASM) criteria^24^ using scalp EEG recordings. Analyses were conducted across three vigilance conditions: N2 sleep, N3 sleep, and a combined NREM category consisting of N2 and N3 epochs. Unless otherwise specified, primary analyses focused on NREM sleep. Consistent with previous slow oscillation studies, REM sleep and stage N1 were excluded from all analyses^8,10^.

### 2.4. SEEG Channel Polarity Determination

In scalp EEG recordings, the orientation of cortical current dipoles relative to the recording electrodes is largely fixed, such that the slow oscillation (SO) down-state (neuronal silence) consistently appears as a negative deflection. In contrast, SEEG contacts may sample local field potentials from different spatial orientations, allowing the physiological down-state to manifest as either a positive or negative deflection depending on electrode geometry and anatomical position^11,25^. If polarity is not corrected, SO trough timing may be misidentified, resulting in phase-assignment errors and potentially misleading estimates of SO-related coupling measures.

To ensure consistent physiological interpretation across channels, SEEG polarity was determined using a gamma-power–based approach previously described in the literature^25^. This method exploits the characteristic relationship between SO phase and gamma activity during NREM sleep, whereby gamma power (30–80 Hz) is reduced during the down-state and increases during the subsequent up-state. For each channel, the 25% largest SO candidates (ranked by absolute trough amplitude) were selected from the NREM sleep event pool. Gamma power was calculated using the Hilbert transform of a 30–80 Hz signal obtained with a 4th-order Butterworth bandpass filter. Mean gamma power was then computed separately for the negative half-wave (from SO onset to the midpoint zero-crossing) and the positive half-wave (from the midpoint zero-crossing to SO termination). The physiological down-state was defined as the half-wave exhibiting lower mean gamma power. Channels in which the negative half-wave showed lower gamma activity were retained in their original orientation. Conversely, when lower gamma power occurred during the positive half-wave, the channel was classified as polarity-inverted and all trough and peak assignments were reversed, ensuring that the SO trough consistently represented the physiological down-state across all SEEG contacts.

### 2.5. Slow Oscillation Detection

SOs were identified independently in scalp EEG and thalamic SEEG recordings using a zero-crossing detection algorithm based on established methodologies^5,8,26,27^. Signals were first bandpass-filtered between 0.1 and 4 Hz with a 4th-order zero-phase Butterworth filter. Candidate SOs were defined as signal segments spanning the first and third zero-crossings of three consecutive zero-crossings, where the first transition was positive-to-negative and the second was negative-to-positive. Candidates were classified as SOs if the negative half-wave duration ranged from 0.25 to 1.0 s and the total event duration ranged from 0.5 to 5.0 s. Events overlapping artifacts or occurring during N1 or REM sleep were excluded. Following detection, an amplitude criterion was applied separately to each channel, retaining only events within the highest 25% of absolute trough amplitudes. An additional kurtosis threshold (waveform kurtosis < 5.0, calculated from the bandpass-filtered SO waveform) was used to remove events with atypical non-Gaussian morphology. For thalamic contacts, the same detection and kurtosis criteria were employed, and the top 25% of events by absolute trough amplitude were retained for each contact based on the NREM sleep event pool. Residual muscle-related artifacts were reduced using the procedure of Brunner et al., which identifies bins in the 26.25–32 Hz range whose amplitudes exceed four times the median of neighboring bins^28^, together with the broadband (4–50 Hz) method of Wang et al., which excludes 5-s segments exceeding six times the global median^29^. Interictal epileptiform discharges (IEDs) were detected using an automated algorithm applied to each SEEG contact, and SO candidates occurring within ±1 s of a detected IED were excluded from subsequent analyses.

### 2.6. Scalp SO Classification by Propagation Pattern

#### 2.6.1. Propagation-Type Classification

To characterize the spatial extent of SOs, each detected event was classified as Global, Frontal, or Local based on its co-occurrence pattern across EEG channels using the established algorithm introduced by Malerba et al. (2019)^8^. This approach relies on unsupervised clustering of binary channel co-occurrence patterns. Following this prior work, we set the number of clusters to k = 3, consistent with the established implementation of this algorithm and corresponding to the point at which further increases in k do not yield qualitatively distinct or more stable spatial patterns. For each SO, a binary co-occurrence matrix was constructed by identifying electrodes whose troughs occurred within a 400-ms window of one another. K-means clustering using Hamming distance (k = 3, 1000 replicates) was applied to these binary co-occurrence vectors. Cluster centroids were mapped to scalp topographies and used to assign subtype labels. Global SOs exhibited widespread bilateral engagement across frontal, central, and posterior regions; Frontal SOs were largely restricted to prefrontal and frontal electrodes; and Local SOs occurred sparsely without a coherent spatial pattern. Among SO types, Global and Frontal SOs typically emerged initially in prefrontal and frontal scalp EEG regions, consistent with the prefrontal predominance of SO initiation. Therefore, this study focused primarily on Frontal and Global SOs as propagating SO subtypes.

#### 2.6.2. Rule-Based Classification

A complementary rule-based classification was applied to categorize the spatial propagation extent of each frontally initiated SO. SOs were grouped according to the most posterior scalp region in which trough co-detection was observed within the 400 ms window. Frontal-Only SOs remained confined to frontal electrodes, Frontal-to-Central (F→C) SOs propagated from frontal to central regions without reaching more posterior regions, Frontal-to-Parietal (F→P) SOs propagated from frontal to parietal regions without reaching occipital regions, and Frontal-to-Occipital (F→O) SOs propagated from frontal to occipital regions within the 400 ms window. Each SO was assigned exclusively to the category corresponding to its maximum posterior propagation extent. Only SOs with frontal initiation were included, ensuring that all four groups shared the same frontal origin. This ordinal extent variable enabled monotonic regression analyses to test whether thalamic properties varied continuously with propagation distance.

### 2.7. Temporal and Phase Coupling Analysis

A subset of frontal and prefrontal scalp EEG channels (F3, F4, F7, F8, F9, F10, Fp1, Fp2, and Fz) was selected to examine the association between thalamic slow oscillation activity and the spatial propagation of cortical SOs, classified using both propagation-type and rule-based approaches. These channels were chosen to sample prefrontal cortical (PFC) activity, consistent with established approaches for investigating frontal SO dynamics^12^. Accordingly, PFC SOs in this study refer to SOs identified in frontal and prefrontal scalp EEG channels.

#### 2.7.1. Peri-Event Histograms

To characterize the temporal co-occurrence of thalamic SOs relative to PFC SO troughs, peri-event histograms were computed for each thalamic nucleus and sleep stage. For each PFC SO trough event (time=0), the co-occurrence probability of thalamic SO troughs within a ±1 s window (50 ms bins, 40 bins total) was computed and normalized by the number of PFC SO reference events. A control distribution was generated from event-free NREM sleep windows, excluding ±3 s buffers around all detected oscillatory events. Group-level observed and control histograms were averaged across contacts per nucleus and stage. Statistical significance was assessed by cluster-based permutation testing (1,000 permutations, α = 0.05, two-tailed). For each permutation, the thalamic SO timestamps were circularly shifted by a random offset relative to the PFC SO reference events, and the permutation-derived histogram was compared to the observed. Clusters of contiguous suprathreshold bins were identified, and the maximum cluster-level statistic from the permutation distribution was used as the null.

#### 2.7.2. Circular Phase-Locking

To quantify the preferred timing of thalamic SOs relative to the PFC SO cycle, we computed phase coupling between thalamic SO trough times and the instantaneous phase of the PFC SO signal. The PFC scalp EEG signal was bandpass filtered between 0.5 and 1.5 Hz using a zero-phase FIR filter, and the Hilbert transform was applied to obtain the instantaneous phase at each time point. The phase of the PFC SO signal at the time of each thalamic SO trough falling within a ±0.5 s window centered on a PFC SO trough was extracted for each thalamic contact and sleep stage. Group-level phase non-uniformity was assessed using the Rayleigh test for circular non-uniformity, yielding a Rayleigh Z statistic and corresponding p-value. Multiple comparison correction across nuclei was performed using the Benjamini–Hochberg false discovery rate (FDR) procedure within each sleep stage. The phase convention throughout is: 0 degrees = PFC SO up-state peak; ±180 degrees = down-state trough; negative angles = ascending post-trough phase (moving from down-state toward up-state); positive angles = descending pre-trough phase. Nuclei with a single recording contact were excluded from inferential phase coupling analysis, as the group MVL = 1 and Rayleigh Z = 1 are mathematical properties of single-contact circular statistics and do not reflect group-level phase consistency.

### 2.8. Thalamic SO Waveform Characteristics and Association with Cortical SO Propagation

To test whether the intrinsic waveform properties of thalamic SOs differed according to the cortical SO propagation pattern with which they co-occurred, we extracted morphological features from thalamic SO events occurring within ±500 ms of a PFC SO trough classified as either Global or Frontal using the propagation-type classification, or within the four propagation categories defined by the rule-based classification. Five waveform features were extracted from each thalamic SO event: (1) SO amplitude (absolute trough amplitude, μV), (2) SO peak-to-peak amplitude (voltage difference between the trough and subsequent positive peak, μV), (3) SO duration (time interval between the first and third zero-crossings, s), (4) descending slope (absolute trough amplitude divided by the time interval from the first zero-crossing to the negative peak; μV/s), and (5) ascending slope (peak-to-peak amplitude divided by the time interval between the negative and positive peaks; μV/s).

To compare Global and Frontal SOs, linear mixed-effects (LME) models were fitted separately for each waveform feature, thalamic nucleus, and sleep stage. SO propagation type was entered as a binary fixed effect with Frontal SO as the reference category, and random intercepts were included for subject and recording contact to account for repeated measurements within participants and within contacts across sessions. The regression coefficient for Global SOs therefore reflected the estimated mean difference in thalamic waveform features between Global and Frontal SO-associated events. The regression coefficient for Global SOs (β) therefore reflected the estimated mean difference between Global and Frontal SO-associated thalamic events. FDR correction was applied across all nucleus × feature comparisons within each sleep stage. To test whether waveform morphology varied continuously with cortical propagation extent, complementary Spearman rank correlations were computed between each waveform feature and the ordinal rule-based propagation extent variable (Frontal-Only, F→C, F→P, F→O). Positive correlation coefficients indicated increasing feature values with greater cortical propagation extent, whereas negative coefficients indicated progressively smaller feature values with wider propagation.

### 2.9 Pre-Onset Thalamic SO Spectral Features

To characterize the thalamic pre-onset state preceding Global versus Frontal PFC SOs, spectral and cross-frequency coupling features were extracted from thalamic SEEG contacts during a variable-length window extending from 2 s before each PFC SO trough to SO onset. A 1-s baseline window from 3 to 2 s before the trough was used for normalization. Three feature categories including spectral and cross-frequency coupling features, were computed for each thalamic contact and each PFC SO event. Spectral power features were computed from the instantaneous Hilbert amplitude squared in seven frequency bands: delta (δ; 0.5–4 Hz), theta (θ; 4–8 Hz), alpha (α; 8–12 Hz), sigma (σ; 12–16 Hz), beta (β; 16–30 Hz), low-gamma (Lγ; 30–59 Hz), and high-gamma (Hγ; 61–119 Hz). Power values were expressed as percent change from baseline.

Phase–amplitude coupling (PAC) features quantified the influence of low-frequency phase on higher-frequency amplitude fluctuations^30^. For each SO event, phases were extracted using Hilbert transforms of band-pass–filtered signals (delta through alpha), and amplitude envelopes were extracted from higher-frequency bands (theta through high gamma). The Modulation Index (MI) provided a normalized measure of coupling strength and was computed as the normalized Kullback–Leibler (KL) divergence of the phase–amplitude distribution from uniformity^31^. MI was calculated by binning the phase of the low-frequency oscillation into N = 18 equally spaced bins (20° per bin) and computing the mean amplitude of the high-frequency envelope within each phase bin to construct a discrete phase–amplitude distribution, P. This distribution was then compared with the uniform distribution U = 1/N using the KL divergence, defined as Equations 1 and 2:

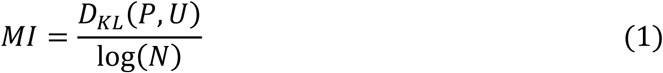

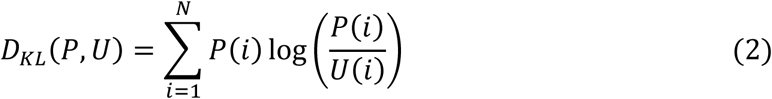

This normalization bounds MI between 0 (no coupling) and 1 (maximal coupling)^31^. The resulting MI values were additionally baseline-normalized by subtracting the MI computed in the matched non-event baseline window. Amplitude-amplitude coupling (AAC) features were computed as Pearson correlations between Hilbert amplitude envelopes for all unique frequency-band pairs and were baseline-subtracted relative to the pre-trough baseline window.

For each feature, thalamic nucleus, and sleep stage, Global and Frontal SOs were compared using the LME model described in Section 2.8. The coefficient for Global SOs therefore represented the estimated difference in pre-onset thalamic activity preceding Global versus Frontal SOs. Random intercepts for Subject and Contact were included to account for repeated measurements within participants and recording contacts. FDR correction was applied within each feature family and sleep stage. Results are reported for features surviving FDR correction.

### 2.10 Statistical Analysis

All analyses were implemented in MATLAB (MathWorks, version 2025b). Circular statistics (Rayleigh test, mean vector length, mean resultant direction) were computed using the CircStat toolbox^32^. For peri-event histogram significance, cluster-based permutation testing (1,000 iterations, α = 0.05, two-tailed) was used. All reported p-values are two-tailed. Statistical significance was defined as p < 0.05 (nominal) or FDR-corrected p < 0.05, as appropriate. Effect size measures included Spearman correlation coefficients (r), mean vector length (MVL), and Rayleigh Z statistics, and Cohen’s d for pairwise between-nucleus comparisons, where applicable. Multiple-comparison correction procedures differed across analyses according to the underlying statistical framework. For peri-event histograms, cluster-based permutation testing inherently corrected across time bins within each histogram^33^. For phase-coupling analyses, Benjamini–Hochberg FDR correction was applied across nuclei within each sleep stage. For waveform morphology analyses, FDR correction was applied across nucleus × feature comparisons within each sleep stage. For pre-onset spectral and cross-frequency features, FDR correction was applied within each feature family and sleep stage. To directly test whether phase-coupling strength differed across nuclei, a linear mixed-effects model was fitted to contact-level MVL values, with nucleus, SO type, and their interaction included as fixed effects and patient included as a random intercept. Analyses were restricted to nuclei represented by at least 6 contacts from at least two patients. Post hoc pairwise comparisons between nuclei were performed with FDR correction. To test whether preferred phase angles differed across nuclei, pairwise permutation tests were conducted (1,000 permutations), with nucleus labels randomly permuted across the pooled contact-level phase distributions.

## 3. Results

### 3.1. Overview of Recording Configuration and Scalp SO Characteristics

We analyzed 24 full-night scalp EEG–SEEG recording sessions from 6 patients across 11 thalamic nuclei. Figure 1A schematically illustrates the recording modalities used to acquire signals from the PFC using prefrontal and frontal scalp EEG channels, and from the thalamus and its nuclei using SEEG depth electrodes. Figure 1B shows a sample SO detected in the Fz channel during N3 sleep. Figure 1C shows that scalp SOs exhibited a clear anterior predominance during NREM sleep, with frontal electrodes displaying the most negative trough amplitudes and highest SO densities, both of which declined progressively along the anteroposterior gradient. This spatial distribution was consistent across N2 and N3 sleep (Supplementary Figure S1) and is in line with the established prefrontal origin of slow oscillatory activity during human NREM sleep^5,26,34^.

**Figure 1.**
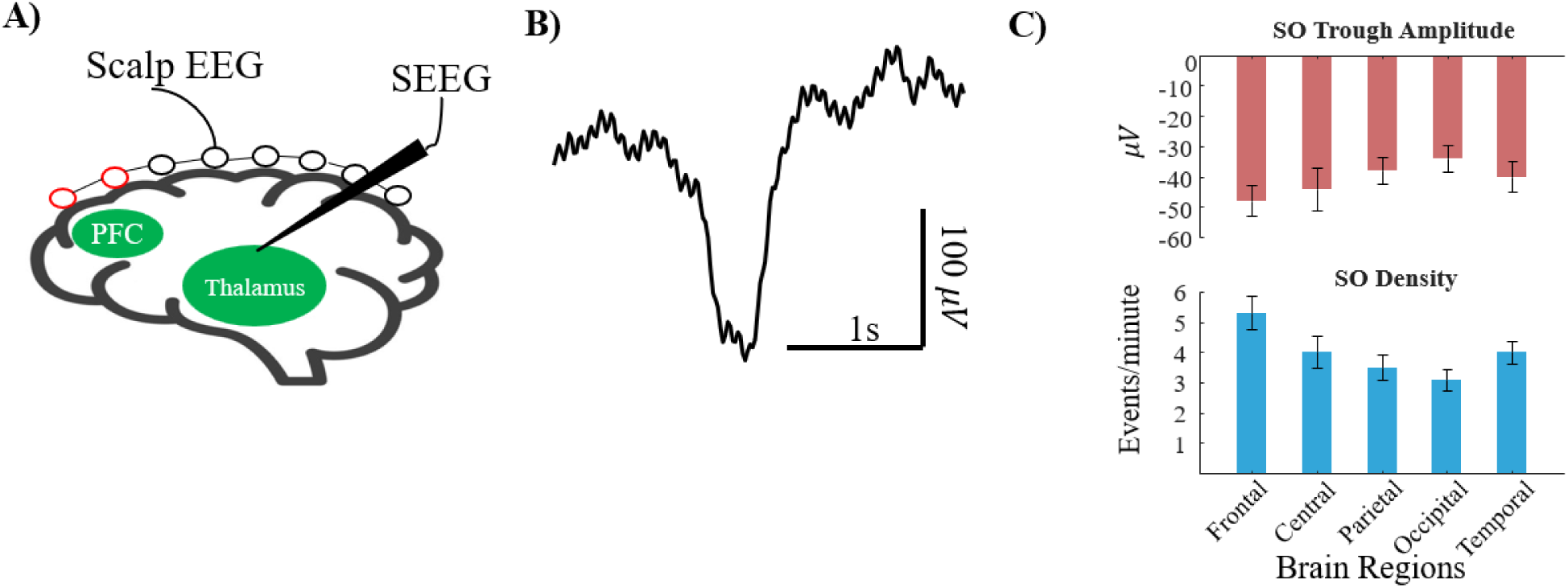
Experimental overview. **A)** Schematic illustrating the simultaneous scalp EEG and SEEG recording configuration used to record cortical SO activity from the scalp and SO activity from multiple thalamic nuclei. Full-coverage scalp EEG channels were used to capture traveling SOs across the cortex, while the red channels indicate prefrontal and frontal scalp electrodes used to record neocortical SO activity from the PFC. Depth SEEG electrodes implanted in 11 distinct thalamic nuclei simultaneously sampled intrathalamic dynamics. **B)** Representative scalp EEG SO waveform recorded from a frontal channel (Fz) during N3 sleep, illustrating the canonical slow oscillation morphology: a large-amplitude negative half-wave (down-state trough) followed by a positive half-wave (up-state peak). **C)** Spatial distribution of scalp SO properties across five scalp regions (Frontal, Central, Parietal, Occipital, Temporal) during NREM sleep, shown as group mean ± SEM SO trough amplitude (μV, upper panel) and SO density (events/min, lower panel). Both measures show a clear frontal predominance, with the most negative trough amplitudes and the highest event rates at frontal electrodes, declining monotonically along the anteroposterior gradient, consistent with the established prefrontal origin of canonical slow oscillations.

### 3.2. Thalamic Nuclei Broadly Co-Activate with neocortical SOs During NREM Sleep

To characterize how individual thalamic nuclei temporally relate to cortical SO activity, we computed peri-event histograms of thalamic SO trough occurrence relative to the PFC SO trough (time = 0; ±1 s window, 50 ms bins) for all nuclei across NREM sleep. All thalamic nuclei with sufficient recording data showed significant temporal co-occurrence with PFC SOs (cluster-based permutation test, p ≤ 0.035 for all; Table 1).

**Table 1.** Temporal and phase coupling between thalamic SOs and PFC SO troughs during NREM sleep across 11 thalamic nuclei and the pooled thalamic signal. Temporal coupling was assessed using peri-event histograms (±1 s window, 50 ms bins): Max Rate (%) = highest group-level co-occurrence probability observed across the analysis window; Peak Bin (s) = time of this maximum rate relative to the PFC SO trough (time = 0; negative = thalamic SO leads PFC trough; positive = thalamic SO lags). Cluster-based permutation testing (1,000 permutations, α = 0.05, two-tailed) was used to assess significance; blue shading indicates nuclei with significant temporal coupling (p < 0.05). Phase coupling was assessed using the Hilbert-transform instantaneous phase of the PFC SO signal (0.5–1.5 Hz bandpass) at the time of each thalamic SO trough falling within ±0.5 s of a PFC SO trough. Group MVL = group mean vector length (0–1; higher values reflect stronger cross-contact phase consistency). Preferred Phase = preferred phase of thalamic SO troughs relative to the PFC SO cycle (0° = PFC SO up-state peak; ±180° = down-state trough; negative values = ascending post-trough phase; positive values = pre-trough descending phase). Rayleigh Z = group-level Rayleigh statistic for circular non-uniformity. Blue shading indicates nuclei with significant phase coupling (Rayleigh p < 0.05; FDR-corrected). Grey shading indicates nuclei with a single recording contact (N = 1), for which MVL = 1.000 and Rayleigh Z = 1.000 are mathematical properties of single-observation statistics and should not be interpreted as evidence of phase coupling.

|  |  | Temporal Coupling |  | Phase Coupling |  |  |
| --- | --- | --- | --- | --- | --- | --- |
| Nuclei | Contact Count | Max Rate (%) | Max Rate Bin (ms) | Group MVL | Preferred Phase ($^\circ$) | Rayleigh Z |
| Pooled all nuclei | 24 | 8.52 | 150 | 0.31 | -157 | 2.25 |
| PuM | 78 | 1.02 | 150 | 0.23 | -148 | 4.16 |
| PuL | 21 | 1.07 | -100 | 0.15 | -176 | 0.46 |
| PuA | 4 | 0.88 | 150 | 0.51 | -38 | 1.05 |
| CM | 16 | 1.24 | 150 | 0.30 | -149 | 1.41 |
| MD | 6 | 1.55 | 150 | 0.68 | 5 | 2.81 |
| VPL | 32 | 1.32 | 150 | 0.16 | -44 | 0.79 |
| VL | 12 | 2.06 | 350 | 0.42 | -98 | 1.39 |
| CL | 6 | 1.53 | -350 | 0.30 | 31 | 0.52 |
| ANT | 1 | 2.62 | -450 | 1 | -162 | 1 |
| VA | 1 | 5.80 | 500 | 1 | 12 | 1 |
| Pf | 4 | 1.49 | 500 | 0.50 | -86 | 1.01 |

Critically, the co-occurrence profiles were characterized by broad, sustained elevations spanning the full ±1 s analysis window, rather than the sharp, isolated peaks reported by Schreiner et al. (2022) for ANT and MD nuclei. Visual inspection of peri-event histograms revealed subtle but consistent timing differences across nuclei: pooled thalamic contacts peaked 150 ms after the PFC SO trough (Figure 2A), PuL peaked 100 ms before the PFC SO trough (Figure 2B), and PuM peaked 150 ms after the trough (Figure 2C).

**Figure 2.**
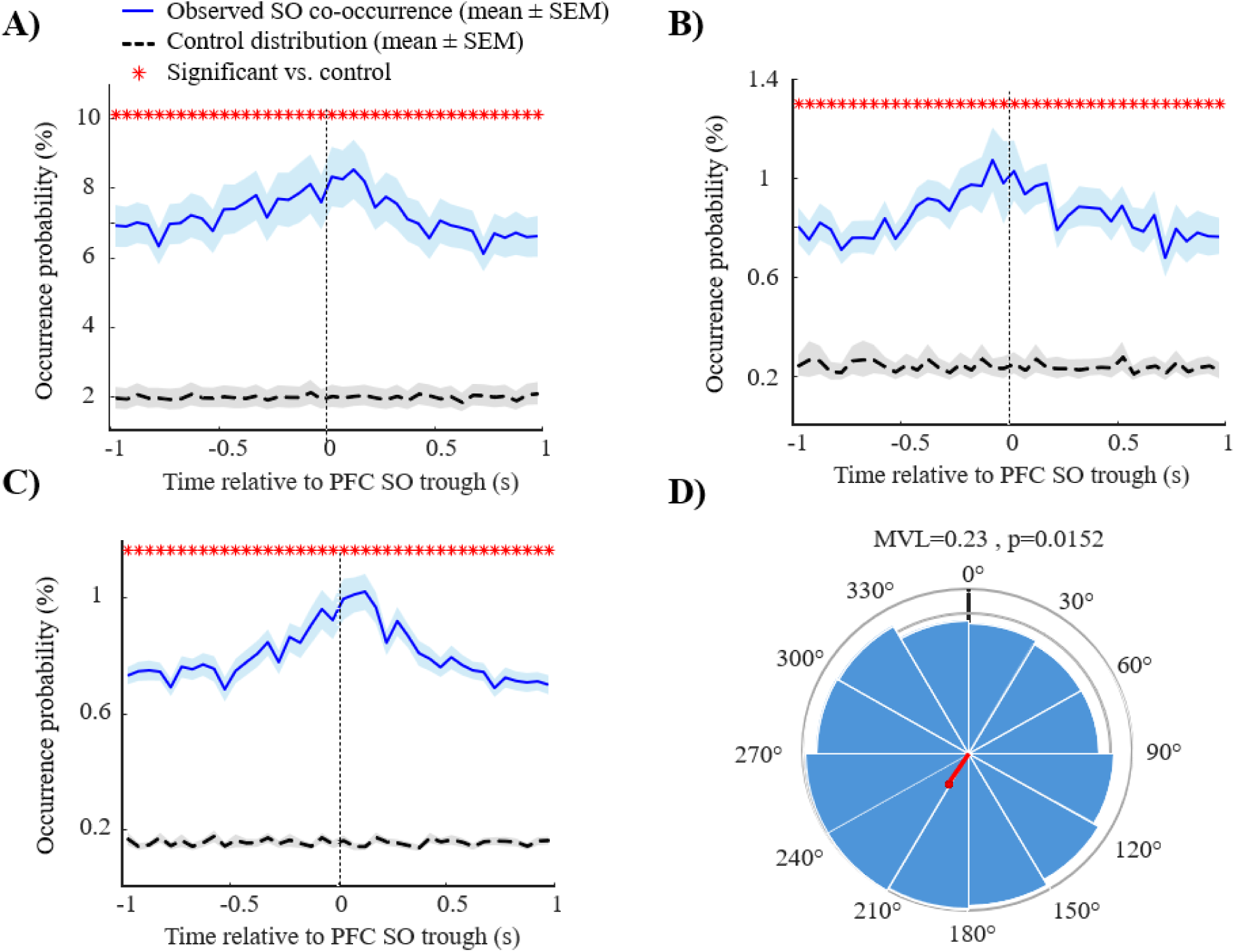
Temporal co-occurrence and phase coupling of thalamic nucleus SOs locked to PFC SO troughs during NREM sleep. Thalamic SOs exhibit broad, sustained co-occurrence with PFC SO troughs across all examined nuclei, with nucleus-specific timing tendencies and phase preferences. **A)** Group-level peri-event histogram (±1 s window, 50 ms bins) for pooled thalamic contacts (N = 24 contacts, NREM), showing the co-occurrence probability of thalamic SO troughs relative to PFC SO trough time (time = 0). The histogram displays a broad, sustained elevation spanning the full window, peaking near 150 ms after the PFC SO trough. Red asterisks mark bins with significant co-occurrence elevation above control (cluster-based permutation test; all 40 bins significant, cluster p < 0.001). Dashed black line shows the control distribution; shading around both the observed and control histograms represents ±SEM. **B)** Peri-event histogram for the lateral pulvinar (PuL; N = 21 contacts, NREM), showing a broad elevation peaking approximately 100 ms before the PFC SO trough; all 40 bins significant (cluster p < 0.001). **C)** Peri-event histogram for the medial pulvinar (PuM; N = 78 contacts, NREM), showing a broad elevation peaking approximately 150 ms after the PFC SO trough; all 40 bins significant (cluster p < 0.001). **D)** Phase polar plot for PuM during NREM sleep (±0.5 s coupling window; p = 0.015). The red arrow indicates the group mean preferred phase of thalamic SO troughs relative to the PFC SO cycle (preferred phase = −148° [equivalent to 212° in the unsigned convention], ascending post-trough phase; Rayleigh Z = 4.16, p = 0.015; N = 78 contacts), reflecting significant phase-locking of PuM SOs to the cortical transition from down-state toward up-state. Phase convention: 0° = PFC SO up-state peak; 180° = down-state trough. Negative phase angles (−1° to −180°) = post-trough ascending phase; positive angles (1° to 180°) = pre-trough descending phase.

Despite broadly similar temporal profiles, individual nuclei exhibited distinct phase preferences relative to the PFC SO cycle. The PuM showed the strongest group-level phase locking during NREM, with a significant preferred phase in the ascending post-trough range (preferred phase = −148°, Rayleigh Z = 4.16, p = 0.015; Figure 2D), indicating that PuM slow oscillations tend to co-occur as the cortex transitions from down-state back toward the up-state. The mediodorsal nucleus (MD) showed a strong directional tendency toward the PFC SO up-state during NREM (preferred phase = 5°, Rayleigh Z = 2.81, p = 0.054), narrowly missing significance despite all 6/6 contacts individually locking to the same phase — a result likely reflecting limited statistical power at N = 6 contacts and consistent with neocortical SOs leading MD activity, as previously reported in humans^12^. All other nuclei did not reach group-level phase coupling significance during NREM sleep, despite widespread individual contact-level locking, reflecting heterogeneous preferred phases that cancel at the group level. Stage-specific results for temporal co-occurrence and phase coupling across all nuclei, separately for N2 and N3, are provided in Supplementary Tables S2 and S3, respectively. Notably, CM and PuM showed significant phase coupling in N3, with preferred phases in the ascending post-trough range (CM: Z = 3.29, p = 0.033, preferred phase = −136°; PuM: Z = 5.45, p = 0.004, preferred phase = −131°), but not N2. MD showed significant coupling in N2 (Z = 2.90, p = 0.048), with a preferred phase near the PFC SO up-state peak (preferred phase = +12°), but not N3. Together these results suggest that the degree and timing of thalamic engagement with cortical SOs varies across sleep depth. Together, these findings indicate that thalamic SO coupling is not homogeneous but reflects structured, functionally organized phase relationships within the thalamocortical sleep network.

### 3.3. Cortical SO Propagation Pattern Differentially Engages Thalamic Nuclei

To examine whether thalamic nuclei couple differently to cortical SOs of varying propagation extent, we classified PFC SOs using the propagation-type classification^34^ into Global SOs (co-detected simultaneously across widespread scalp electrodes including frontal, central, and parietal regions) and Frontal SOs (co-detected predominantly at frontal electrodes, reflecting spatially restricted propagation). Local SOs, which occur sparsely across brain regions without a clear spatial pattern, were identified but excluded from the primary comparison. Since both Global and Frontal SOs initially emerge in the frontal scalp region — consistent with the established prefrontal cortical origin of slow oscillations^5,35^ — we examined how each propagation type is associated with distinct thalamic engagement profiles. This distinction is particularly important given that Global SOs have been linked to episodic memory consolidation, a role absent for Frontal SOs^9^, making thalamic differentiation between these types directly relevant to understanding the neural mechanisms of sleep-dependent memory.

Peri-event histograms for both Global and Frontal SOs revealed thalamic co-occurrence across the full ±1 s window in all major nuclei. Group-level histograms revealed subtle timing differences between the two SO types: for pooled thalamic contacts, Global SOs peaked approximately 50 ms after the PFC SO trough while Frontal SOs peaked approximately 150 ms after the trough (Figure 3A). For PuL, Frontal SOs peaked 150 ms before the trough, whereas Global SOs showed a broad, flat elevation without a clear peak, spanning approximately −100 to +50 ms relative to the trough (Figure 3B). For PuM, both Global and Frontal SOs peaked 150 ms after the trough (Figure 3C). The fundamental temporal coupling between thalamus and cortex during slow oscillations is thus largely preserved regardless of propagation extent, with the principal differences emerging at the level of phase organization.

**Figure 3.**
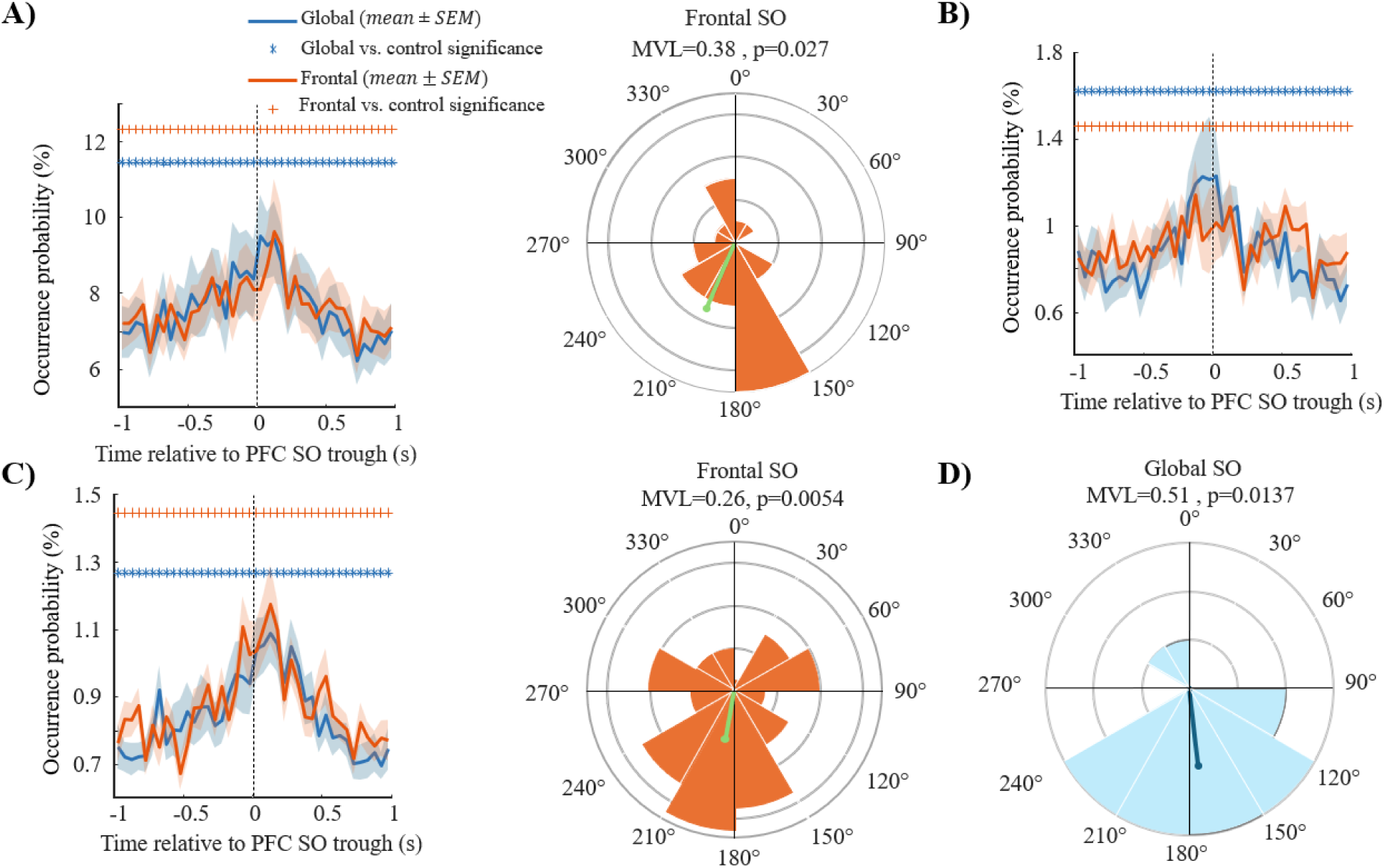
Differential thalamic coupling to Global versus Frontal PFC slow oscillations during NREM sleep. **A)** Overlaid peri-event histograms for pooled thalamic contacts (N = 24 contacts) showing co-occurrence probability for Global (blue) and Frontal (orange) SO types. Both SO types showed elevated thalamic co-occurrence within the ±1 s window surrounding the PFC SO trough. Visual inspection showed that Global SOs peaked 50 ms after the PFC SO trough, whereas Frontal SOs peaked 150 ms after the trough. The polar plot for Frontal SOs (orange) showed significant group-level phase locking during NREM (Rayleigh Z = 3.54, p = 0.027, preferred phase = −156°, ascending post-trough), whereas Global SOs did not reach significance (p = 0.218). **B)** Overlaid peri-event histograms for the lateral pulvinar (PuL; N = 21 contacts). Frontal SOs peaked approximately 150 ms before the PFC SO trough, whereas Global SOs showed a broad, flat elevation without a clear peak, spanning approximately −100 to +50 ms relative to the trough. Neither SO type reached significant group-level phase coupling in PuL during NREM. **C)** Overlaid peri-event histograms and phase polar plots for the medial pulvinar (PuM; N = 78 contacts). Both Global and Frontal SOs peaked 150 ms after the PFC SO trough. The polar plot for Frontal SOs (orange) demonstrated significant phase locking in PuM during NREM (Rayleigh Z = 5.17, p = 0.005, preferred phase = −170°, ascending post-trough), whereas Global SOs did not reach significance. **D)** Phase coupling of the centromedian nucleus (CM) during Global SOs. CM was the only nucleus exhibiting significant phase locking to Global SOs in NREM sleep (Rayleigh Z = 4.14, p = 0.013, preferred phase = 175°), corresponding to the cortical down-state trough.

Phase coupling analyses revealed a striking dissociation between the two SO propagation types during NREM sleep. Frontal SOs drove stronger group-level phase locking in PuM and pooled thalamic contacts compared to Global SOs (Supplementary Tables S4–S5), whereas Global SOs uniquely and robustly synchronized the CM near the cortical down-state trough (preferred phase = 175°, Z = 4.14, p = 0.013; Supplementary Table S5). Specifically, Frontal SOs generated significant phase locking in pooled thalamic contacts during NREM (Rayleigh Z = 3.54, p = 0.027), with a preferred phase in the ascending post-trough range (preferred phase =−156°; Figure 3A polar plot). Among all SO propagation type comparisons, the strongest phase coupling effect was observed in PuM, where Frontal SOs produced the strongest phase locking in the dataset during NREM (Z = 5.17, p = 0.005; preferred phase = −170°; Figure 3C polar plot). In contrast, Global SOs did not reach significance in any nucleus during NREM except CM, which uniquely synchronized near the cortical down-state trough (preferred phase = 175°, Z = 4.14, FDR p = 0.013; Figure 3D). The MD showed a directional tendency toward up-state phase preference for both types (Global: 5.5°; Frontal: 30.6°) without reaching significance in either case during NREM, consistent with neocortical SOs leading MD activity as previously reported^12^ (Supplementary Tables S4–S5). Stage-specific phase coupling results for Global and Frontal SOs separately across N2 and N3 are provided in Supplementary Tables S6 and S7, respectively. In N3, Frontal SOs drove significant phase locking in pooled thalamus (Z = 7.38, p < 0.001), PuM (Z = 9.54, p < 0.001), and PuL (Z = 4.26, p = 0.012). Global SOs reached significance in PuM during N3 (Z = 4.19, p = 0.015, preferred phase = −143°) and CM during both N2 (Z = 3.05, p = 0.045) and N3 (Z = 4.76, p = 0.006), confirming that CM down-state coupling and PuM post-trough coupling are not confined to pooled NREM but are present across individual sleep stages.

To evaluate whether these differences reflect nucleus-specific effects rather than patient composition, we examined recording overlap across nuclei (Supplementary Table S1). PuM was recorded in 5 of 6 patients (between 4 and 24 contacts per patient, 78 contacts total) and CM in 2 of 6 patients (8 contacts per patient, 16 contacts total); these two nuclei shared 2 patients contributing 8 recording nights. For the CM Global SO finding, both patients individually showed phases near the group preferred phase of 175° (patient 003: phase = 158°, Z = 1.91; patient 023: phase = −170°, Z = 2.58; angular deviations of 17° and 15° respectively), confirming the finding is not driven by patient sampling. For the PuM Frontal SO finding, individual analyses across all 5 patients showed preferred phases spanning the full circle (−168° to 167°; Rayleigh Z range: 0.30–8.83), with three of four patients contributing more than 4 contacts showing phases clustered near the group preferred phase of −170° (within 45°), and patient 001 reaching individual significance (Z = 8.83, p < 0.001, preferred phase = −153°, N = 24 contacts), indicating that the phase direction is consistent across the majority of patients.

Direct between-nucleus comparisons across well-sampled nuclei (PuM, CM, PuL, and VPL; ≥6 contacts from ≥2 patients; 246 contacts total; Supplementary Table S1) showed no significant interaction between nucleus and SO type on coupling strength (F(3,238) = 0.24, p = 0.87). Pairwise post-hoc comparisons, focused on the primary comparison of interest (PuM and CM), revealed significantly higher MVL in PuM than CM for both Frontal SOs (mean MVL 0.199 vs. 0.177, Cohen’s d = 0.19, p < 0.001, FDR-corrected) and Global SOs (0.178 vs. 0.157, d = 0.20, p < 0.001), though effect sizes were small. Permutation-based pairwise tests found no significant difference in preferred phase angles between any nucleus pair for either SO type (all p > 0.49), with PuM and CM preferred phases differing by only 7° for Frontal SOs and 34° for Global SOs. Notably, the higher per-contact MVL in PuM for Global SOs did not translate into group-level Rayleigh significance (Z = 1.29, p = 0.28), because individual PuM contacts showed heterogeneous preferred phases that cancelled at the group level; in contrast, the 16 CM contacts showed consistently aligned preferred phases near the down-state trough, producing robust group-level coherence despite lower per-contact MVL. Together, these results indicate that the PuM and CM findings reflect differences in within-nucleus phase locking significance rather than statistically distinguishable differences in coupling strength or phase direction between nuclei, likely reflecting limited statistical power given the small number of CM contacts (N = 16, 2 patients).

### 3.4. Thalamic SO Waveform Morphology Associates with Cortical Propagation Extent

Beyond the timing and phase of thalamic co-activation, we examined whether the intrinsic morphological properties of thalamic SOs — amplitude, duration, and slope — are associated with cortical SO propagation type. This analysis revealed nucleus-specific patterns that differed in direction across thalamic nuclei. Figure 4A illustrates the amplitude difference in PuL: Frontal SOs co-occurred with significantly larger peak-to-peak amplitudes than Global SOs (LME: β = −1.54 μV, FDR p = 0.002; Spearman r = −0.103, FDR p < 0.001). Figure 4B illustrates the opposing pattern in VPL, where Global SOs co-occurred with significantly larger peak-to-peak amplitudes than Frontal SOs (LME: β = 3.18 μV, FDR p < 0.001; Spearman r = 0.094, FDR p = 0.001). This directional reversal between PuL and VPL reflects a broader nucleus-specific dissociation: PuM and PuL consistently showed larger thalamic SO amplitude and longer duration during Frontal SOs, whereas VPL, CL, and CM showed the opposite, with larger amplitude during Global SOs. For PuL specifically, Figure 4C confirms that Frontal SOs also co-occurred with significantly longer thalamic SO durations (LME: β = −0.013 s, FDR p = 0.009; Spearman r = −0.085, FDR p < 0.001). For PuM, Global SOs similarly co-occurred with shorter-duration thalamic events (LME: β = −0.011 s, FDR p = 0.007). For pooled thalamic contacts, Global SOs co-occurred with lower-amplitude events (β = −0.57 μV, FDR p = 0.016). Among VPL, CL, and CM, CL showed β = 4.11 μV (FDR p = 0.004), and CM showed a directional trend in the expected direction, though the LME contrast did not reach significance (β = 0.74 μV, p = 0.166) and the Spearman correlation, while surviving FDR correction, reflected a negligible effect size (r = 0.025, FDR p < 0.05).

**Figure 4.**
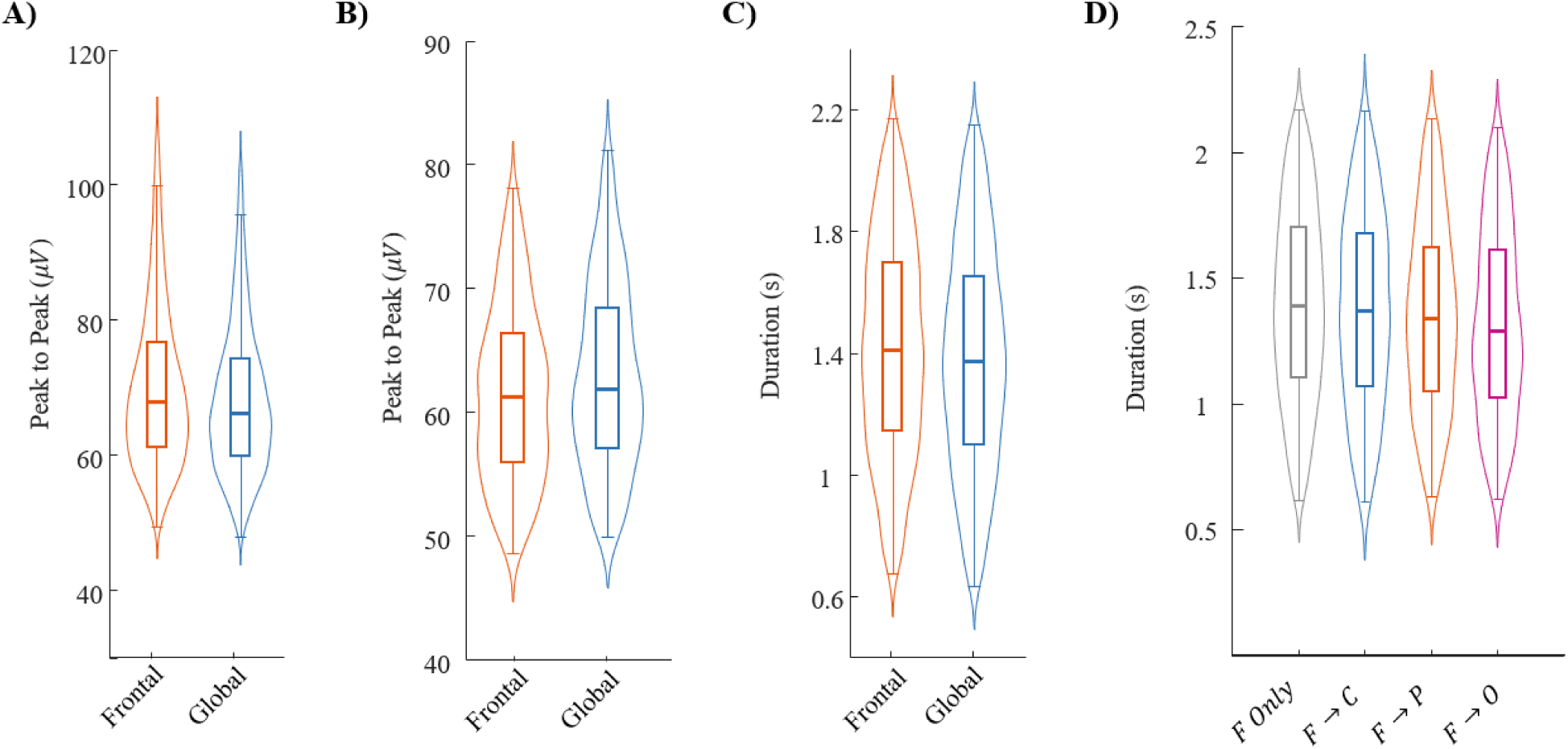
Thalamic SO waveform morphology associates with neocortical SO propagation extent during NREM sleep. A) Violin plots showing peak-to-peak amplitude (μV) of PuL thalamic SO events co-occurring with Frontal (orange) and Global (blue) PFC SOs during NREM sleep. Frontal SOs co-occur with significantly larger peak-to-peak amplitudes in PuL (LME: β = −1.54 μV, FDR p = 0.002; Spearman r = −0.103, FDR p < 0.001). B) Violin plots showing peak-to-peak amplitude (μV) of VPL thalamic SO events co-occurring with Frontal and Global PFC SOs during NREM sleep. Global SOs co-occur with larger peak-to-peak amplitudes in VPL (LME: β = 3.18 μV, FDR p < 0.001; Spearman r = 0.094, FDR p = 0.001), illustrating the opposing directional pattern relative to PuL. C) Violin plots showing duration (s) of PuL thalamic SO events co-occurring with Frontal and Global PFC SOs during NREM sleep. Frontal SOs co-occur with significantly longer duration events in PuL (LME: β = −0.013 s, FDR p = 0.009; Spearman r = −0.085, FDR p < 0.001). D) Violin plots showing that PuL SO duration is associated with the rule-based anteroposterior propagation group (Frontal-Only, F→C, F→P, F→O) during NREM sleep, illustrating a monotonic decrease in PuL thalamic SO duration with increasing propagation extent (Spearman r = −0.085, FDR p < 0.001).

To examine whether these effects scale continuously with propagation extent, we applied the rule-based classification that categorized frontally initiated cortical SOs into four groups by maximal anteroposterior extent: Frontal-Only, F→C, F→P, and F→O. Figure 4D shows that in PuL, thalamic SO duration decreased monotonically with increasing propagation extent (Spearman r = −0.085, FDR p < 0.001), consistent with the Frontal versus Global comparison in Figure 4C. MD and VA also showed progressively increasing amplitude with wider propagation (VA: r = 0.154, FDR p < 0.05; MD: r = 0.056, FDR p < 0.05). For pooled thalamic contacts, SO duration increased monotonically with propagation extent (r = 0.045, FDR p < 0.001).

### 3.5. Pre-Onset Thalamic Dynamics Predict Cortical SO Propagation Type

The finding that thalamic nuclei respond differently to Global versus Frontal SOs raises a critical question with direct implications for neuromodulation: can the thalamus be read out before the cortical SO occurs to predict its upcoming propagation pattern? To address this, we analyzed contact-level thalamic SEEG signals during the 2-second window preceding the neocortical SO trough (corresponding, on average, to a 1.57 s window before SO onset in this dataset; Figure 5A) and examined whether spectral power, phase-amplitude coupling (PAC), and amplitude-amplitude coupling (AAC) in this pre-onset window differed between future Global and Frontal SOs during NREM sleep. Results are reported for PuM, which was the only nucleus in which pre-onset differences between Global and Frontal SOs survived FDR correction (FDR correction within feature type × stage).

**Figure 5.**
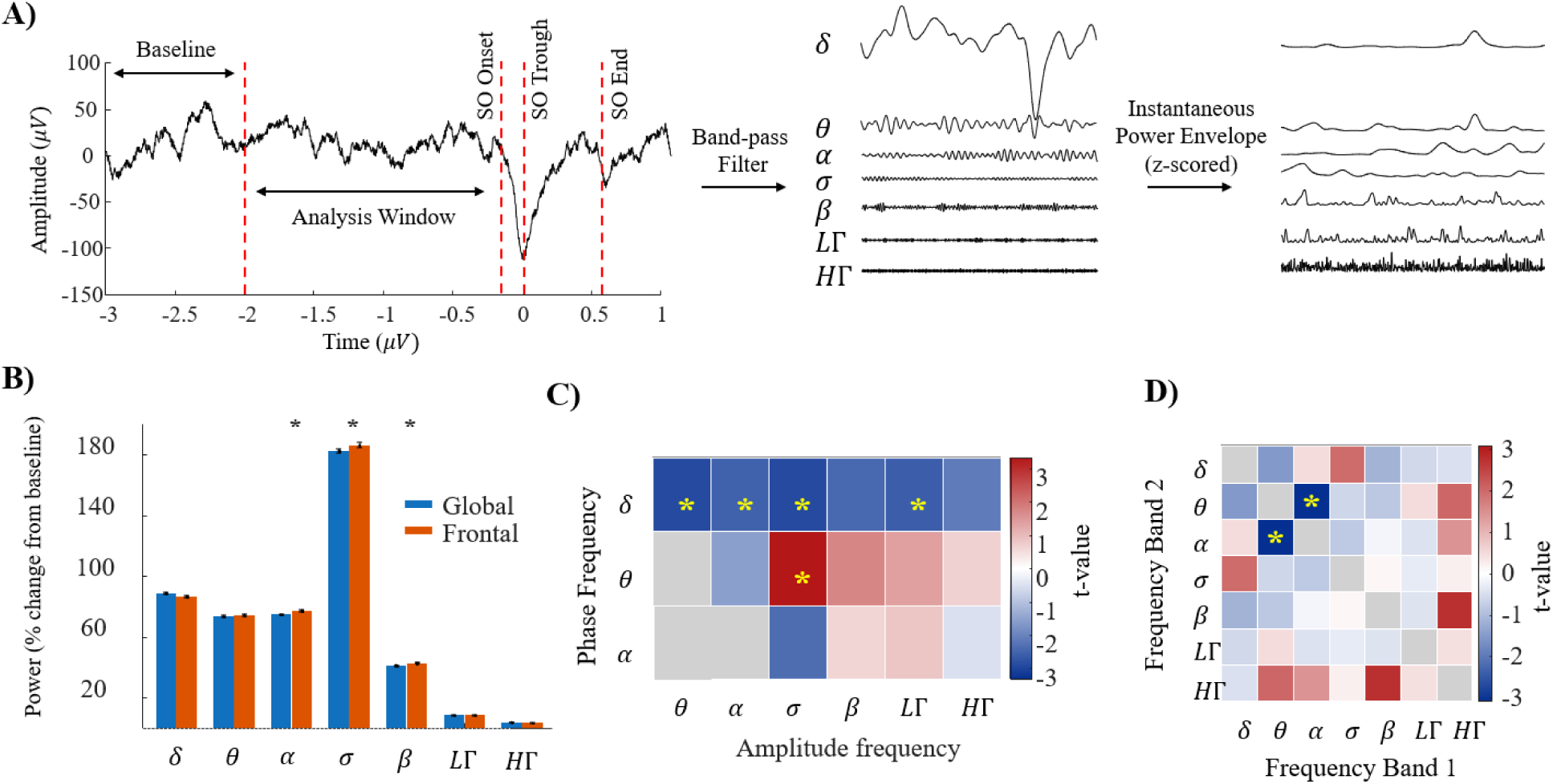
Pre-onset thalamic dynamics in medial pulvinar predict neocortical SO spatial extent. **A)** Schematic of the pre-onset feature analysis pipeline. The panel shows an EEG waveform from 3 s before to 1 s after the PFC SO trough, with band-pass filtered waveforms across seven frequency bands (delta 0.5–4 Hz through high-gamma 61–119 Hz) and their corresponding instantaneous power envelopes (z-scored). Vertical dashed lines demarcate the baseline window (3–2 s before trough), analysis window (2 s before trough to SO onset; mean duration = 1.57 s), SO onset, SO trough (time = 0), and SO end. This panel is adapted from our prior work^22^. **B)** Instantaneous power envelopes across frequency bands in the pre-onset analysis window for PuM thalamic contacts preceding Global (blue) and Frontal (orange) SOs during NREM sleep. Global SOs are preceded by lower alpha-band, sigma-band, and beta-band power compared to Frontal SOs (all FDR p < 0.05). **C)** Phase-amplitude coupling contrast matrix (Global minus Frontal t-values) showing the modulation index between the phase of low-frequency bands (delta, theta, alpha; y-axis) and the amplitude of higher-frequency bands (theta through high-gamma; x-axis), computed during the pre-onset window in PuM during pooled NREM sleep. Global SOs are preceded by significantly reduced delta-phase PAC across all six amplitude bands (all FDR p < 0.0001). In contrast, theta-phase PAC of sigma-band amplitude was elevated before Global SOs (FDR p < 0.0001), indicating a concurrent reorganization of cross-frequency coupling in the pre-onset window. **D)** Amplitude-amplitude coupling correlation matrix showing Pearson r between instantaneous amplitude envelopes across frequency band pairs during the pre-onset window in PuM. Theta–alpha AAC is significantly reduced ahead of Global SOs during NREM (FDR p = 0.044), indicating less coordinated theta–alpha envelope coupling in PuM before widely propagating SOs.

#### Spectral Power

Global SOs were preceded by significantly lower alpha (8–12 Hz), sigma, and beta power in PuM during NREM (all FDR p < 0.05; Figure 5B). These findings indicate that the PuM enters a state of reduced mid-frequency synchronization in the seconds ahead of Global SOs.

#### Phase-Amplitude Coupling

PAC analyses revealed that during NREM, Global SOs were preceded by significantly reduced delta-phase PAC across all six amplitude bands in PuM (all FDR p < 0.0001). The direction reversed for theta-phase PAC of sigma-band amplitude, which was higher before Global SOs (FDR p < 0.0001). These findings indicate that the pre-onset PuM network state is characterized by suppressed delta-phase cross-frequency coupling and a concurrent reorganization of theta-phase coupling ahead of Global SOs (Figure 5C).

#### Amplitude-Amplitude Coupling

AAC analyses revealed that theta–alpha amplitude covariation in PuM was significantly reduced ahead of Global SOs during NREM (FDR p = 0.044; Figure 5D), indicating that less coordinated theta–alpha envelope coupling in the PuM characterizes the pre-onset window before widely propagating SOs.

Collectively, these pre-onset features demonstrate that PuM enters quantifiably distinct network states in the seconds preceding Global versus Frontal SOs, with suppressed delta-phase PAC across amplitude frequencies representing the most spatially consistent discriminating feature.

## 4. Discussion

This study provides a multi-nucleus characterization of thalamocortical SO coupling during human NREM sleep using simultaneous scalp EEG and thalamic SEEG recordings across 11 thalamic nuclei. This recording configuration allowed direct examination of how individual thalamic nuclei relate to cortical slow oscillations within the same events. Several principal findings emerge. First, thalamic SO activity showed broad temporal synchronization with prefrontal cortical SOs across a diverse set of thalamic nuclei during NREM sleep. Second, despite this widespread synchrony, distinct thalamic nuclei occupied different phase positions within the cortical SO cycle, indicating substantial temporal heterogeneity within the thalamus during sleep. Third, thalamic dynamics varied systematically with cortical SO propagation extent, including nucleus-specific differences in phase organization and waveform morphology. Finally, pre-onset activity in the medial pulvinar differed before Global and Frontal SOs, suggesting that thalamic network state prior to SO onset may relate to the subsequent spatial propagation pattern of cortical SOs. Together, these findings extend prior work demonstrating thalamic involvement in sleep slow oscillations and provide evidence that thalamic SO dynamics are spatially and temporally differentiated across nuclei. Rather than behaving as a functionally uniform structure during NREM sleep, the thalamus appears to participate in SO activity through anatomically heterogeneous coupling patterns that vary according to both thalamic nucleus identity and cortical SO propagation structure. These observations contribute to current models of thalamocortical sleep organization and may help inform future studies investigating the role of thalamic networks in sleep-dependent cognition and neuromodulation.

The nucleus-specific phase preferences observed here are worth examining in detail. The MD preferentially aligned near the cortical SO up-state, consistent with prior human intracranial recordings showing that MD slow oscillations follow neocortical SO activity^12^. In contrast, PuM showed robust coupling to the ascending post-trough phase across NREM sleep, while CM showed a similar directional preference that reached significance specifically during N3, suggesting engagement of both nuclei as the cortex transitions from down-state back toward the up-state. The post-trough ascending preference of PuM is also consistent with prior human intracranial recordings showing that pulvinar SOs tend to follow neocortical SO activity^36^. Notably, pooled thalamic analyses showed relatively weak overall phase coherence despite widespread contact-level locking, indicating that averaging across nuclei obscures the structured nucleus-specific timing relationships present within the thalamocortical SO network. In the present dataset, PuM and CM exhibited stronger phase organization during N3 sleep, whereas MD coupling was more prominent during N2, suggesting that the timing and strength of thalamocortical SO coordination may vary across both thalamic nucleus identity and sleep depth, likely reflecting differences in thalamocortical connectivity and intrinsic thalamic physiology.

The present findings further demonstrate that thalamic SO organization varies according to the spatial propagation extent of cortical slow oscillations. Although both Global and Frontal SOs produced broadly similar temporal co-occurrence profiles across thalamic nuclei, differences emerged in phase organization and waveform morphology, indicating that cortical propagation structure is associated with differences in how individual thalamic nuclei participate in SO dynamics. Frontal SOs were associated with stronger and more coherent phase locking in PuM, whereas Global SOs showed more coherent phase synchronization with CM near the cortical down-state. These findings suggest that distinct cortical SO propagation patterns engage partially different thalamocortical networks rather than simply reflecting stronger or weaker versions of the same oscillatory process. Importantly, the direction of these effects varied across nuclei, supporting the idea that thalamic participation during sleep depends not only on SO occurrence, but also on the spatial organization of cortical recruitment.

This CM–Global SO dissociation is of particular functional interest. The CM is an intralaminar thalamic nucleus with widespread cortical and subcortical connectivity, including projections to prefrontal and sensorimotor regions, and has been implicated in cognitive, sensory, and motor functions, supporting a potential role in large-scale thalamocortical coordination during sleep^37^. Given that Global SOs — but not Frontal SOs — are associated with episodic memory consolidation and exhibit stronger sleep spindle coupling^9^, the preferential engagement of CM during Global SO down-states may reflect a thalamocortical mechanism supporting the broad cortical synchronization required for memory-relevant SO propagation. Whether CM activity plays a causal role in enabling the widespread cortical recruitment that characterizes Global SOs, or instead reflects engagement concurrent with the propagating cortical wave, could be addressed in future studies combining intracranial recordings with closed-loop thalamic stimulation. Direct between-nucleus statistical comparisons did not reveal a significant interaction between nucleus and SO type on coupling strength, likely reflecting limited power given the small number of CM contacts, and these findings should therefore be interpreted as within-nucleus effects rather than statistically confirmed nucleus-level dissociations.

The rule-based propagation analysis further demonstrated that thalamic waveform morphology changes systematically with increasing cortical propagation extent. In PuM and PuL, wider cortical propagation was associated with reduced SO amplitude and duration, whereas VPL, CL, and CM showed the opposite trend, with larger thalamic SO amplitudes accompanying broader cortical propagation. These propagation-related gradients were consistent across sleep stages, suggesting that thalamic nuclei differ in how they are engaged within thalamocortical circuits during large-scale SO propagation. Together, these findings indicate that cortical SO propagation is associated with nucleus-specific reconfiguration of thalamic dynamics. Rather than showing uniform coupling patterns across SO propagation types, different thalamic nuclei appear to participate in SO propagation through distinct temporal and morphological patterns, likely reflecting differences in connectivity and network function across thalamocortical circuits.

Thalamic activity preceding cortical SO onset differed according to the subsequent propagation pattern of the cortical SO. In particular, PuM exhibited distinct pre-onset spectral and cross-frequency coupling features before Global versus Frontal SOs, suggesting that thalamic network state prior to SO onset is associated with the later spatial organization of cortical slow oscillatory activity. The pattern of reduced mid-frequency power and suppressed delta-phase cross-frequency coupling in PuM before Global SOs suggests a state of reduced thalamic synchronization preceding widespread cortical recruitment, potentially reflecting a permissive network state that allows broader cortical engagement. The concurrent reorganization of theta-phase coupling further suggests that pre-onset thalamic dynamics are not simply a uniform suppression of activity but involve a reconfiguration of cross-frequency interactions that may differentially gate the spatial extent of subsequent cortical SO propagation.

These findings complement our recent scalp EEG work demonstrating that pre-onset cortical activity contains information about upcoming SO propagation patterns^22^, and extend this concept to the thalamus, suggesting that network signatures associated with SO propagation extent are not restricted to cortical recordings alone. However, because the present analyses are correlational, the findings do not establish whether thalamic pre-onset dynamics actively contribute to determining SO propagation extent or instead reflect broader network states that co-emerge across cortex and thalamus.

Although preliminary given the limited number of subjects and contacts contributing to the pre-onset analysis, these results may have implications for future closed-loop neuromodulation approaches targeting sleep oscillations. Current SO stimulation systems typically operate reactively after SO detection^21^, limiting temporal precision. The present findings demonstrate that PuM pre-onset network states differ before Global versus Frontal SOs, providing proof-of-concept that thalamic activity may contain propagation-relevant information before cortical SO emergence. Future studies with larger cohorts and prospective closed-loop designs will be needed to determine whether these signatures are sufficiently reliable and generalizable to inform real-time stimulation targeting. PuM — with its widespread cortical connectivity and the pre-onset sensitivity observed here — represents a candidate recording site for such predictive closed-loop systems.

Several limitations should be considered when interpreting the present findings. First, the study cohort consisted of patients with drug-resistant focal epilepsy undergoing presurgical SEEG monitoring. Although seizure-associated periods were excluded from analysis, underlying pathology and anti-seizure medications may influence thalamocortical oscillatory dynamics relative to healthy populations. Second, thalamic electrode placement was determined by clinical considerations, resulting in unequal sampling across nuclei and preferential sampling of nuclei with high connectivity to epileptogenic brain regions. Some nuclei, including PuM and VPL, were relatively well represented, whereas others such as ANT and VA had only a single recording contact and could not support group-level inference; additionally, not all nuclei were recorded in all patients, with PuM and CM sharing only 2 of 6 patients. Findings from sparsely sampled nuclei should therefore be interpreted as descriptive rather than inferential. Third, the pre-onset analyses were restricted to PuM. Although pre-onset features survived FDR correction, the limited subject count means these findings should be considered preliminary and require replication in larger cohorts with broader thalamic coverage. Finally, the study is observational and correlational in nature. While the findings demonstrate structured relationships between cortical SO propagation and thalamic dynamics, they do not establish causal roles for specific thalamic nuclei in SO generation, propagation, or coordination. Future studies combining intracranial recordings with causal perturbation approaches, such as closed-loop stimulation, will be necessary to directly test these mechanisms.

This study demonstrates that thalamocortical slow oscillation dynamics during human NREM sleep are both nucleus-specific and associated with cortical SO propagation structure. Although thalamic nuclei broadly synchronize with prefrontal cortical SOs, individual nuclei occupy distinct temporal positions within the cortical SO cycle and exhibit different patterns of engagement during Global versus Frontal SO propagation. In addition, thalamic waveform morphology and pre-onset network activity are systematically associated with cortical propagation extent, particularly within the medial pulvinar for pre-onset network signatures and the centromedian nucleus for propagation-specific phase coupling. Together, these findings support a model in which the human thalamus participates in sleep slow oscillations through nucleus-specific and dynamically organized thalamocortical interactions rather than as a functionally uniform structure. These results extend current understanding of human thalamocortical SO organization and provide a foundation for future studies investigating the role of thalamic networks in sleep-dependent cognition and closed-loop neuromodulation strategies targeting specific SO states.

## References

1. Staresina, B. P., Niediek, J., Borger, V., Surges, R. & Mormann, F. How coupled slow oscillations, spindles and ripples coordinate neuronal processing and communication during human sleep. Nature Neuroscience 26, 1429–1437 (2023).

2. Staresina, B. P. Coupled sleep rhythms for memory consolidation. Trends in Cognitive Sciences (2024).

3. Timofeev, I. & Chauvette, S. Sleep slow oscillation and plasticity. Current opinion in neurobiology 44, 116–126 (2017).

4. Steriade, M., McCormick, D. A. & Sejnowski, T. J. Thalamocortical oscillations in the sleeping and aroused brain. Science 262, 679–685 (1993).

5. Massimini, M., Huber, R., Ferrarelli, F., Hill, S. & Tononi, G. The sleep slow oscillation as a traveling wave. Journal of Neuroscience 24, 6862–6870 (2004).

6. Staresina, B. P. et al. Hierarchical nesting of slow oscillations, spindles and ripples in the human hippocampus during sleep. Nature neuroscience 18, 1679–1686 (2015).

7. Helfrich, R. F. et al. Bidirectional prefrontal-hippocampal dynamics organize information transfer during sleep in humans. Nature communications 10, 3572 (2019).

8. Malerba, P., Whitehurst, L. N., Simons, S. B. & Mednick, S. C. Spatio-temporal structure of sleep slow oscillations on the electrode manifold and its relation to spindles. Sleep 42, (2019).

9. Niknazar, H., Malerba, P. & Mednick, S. C. Slow oscillations promote long-range effective communication: The key for memory consolidation in a broken-down network. Proceedings of the National Academy of Sciences 119, e2122515119 (2022).

10. Alipour, M. et al. The Space–Time Organisation of Sleep Slow Oscillations as Potential Biomarker for Hypersomnolence. Journal of Sleep Research e70059 (2025).

11. Frauscher, B. et al. Facilitation of epileptic activity during sleep is mediated by high amplitude slow waves. Brain 138, 1629–1641 (2015).

12. Schreiner, T., Kaufmann, E., Noachtar, S., Mehrkens, J.-H. & Staudigl, T. The human thalamus orchestrates neocortical oscillations during NREM sleep. Nature communications 13, 5231 (2022).

13. Hughes, S. W., Cope, D. W., Toth, T. I., Williams, S. R. & Crunelli, V. All thalamocortical neurones possess a T-type Ca2+ ‘window’current that enables the expression of bistability-mediated activities. The Journal of physiology 517, 805–815 (1999).

14. McCORMICK, D. A. & Pape, H.-C. Properties of a hyperpolarization-activated cation current and its role in rhythmic oscillation in thalamic relay neurones. The Journal of physiology 431, 291–318 (1990).

15. Blethyn, K. L., Hughes, S. W., Tóth, T. I., Cope, D. W. & Crunelli, V. Neuronal basis of the slow (< 1 Hz) oscillation in neurons of the nucleus reticularis thalami in vitro. Journal of Neuroscience 26, 2474–2486 (2006).

16. Crunelli, V. & Hughes, S. W. The slow (< 1 Hz) rhythm of non-REM sleep: a dialogue between three cardinal oscillators. Nature neuroscience 13, 9–17 (2010).

17. Gent, T. C., Bandarabadi, M., Herrera, C. G. & Adamantidis, A. R. Thalamic dual control of sleep and wakefulness. Nature neuroscience 21, 974–984 (2018).

18. Hay, Y. A. et al. Thalamus mediates neocortical Down state transition via GABAB-receptor-targeting interneurons. Neuron 109, 2682–2690 (2021).

19. Urbain, N., Fourcaud-Trocmé, N., Laheux, S., Salin, P. A. & Gentet, L. J. Brain-state-dependent modulation of neuronal firing and membrane potential dynamics in the somatosensory thalamus during natural sleep. Cell reports 26, 1443–1457 (2019).

20. Slézia, A., Hangya, B., Ulbert, I. & Acsády, L. Phase advancement and nucleus-specific timing of thalamocortical activity during slow cortical oscillation. Journal of neuroscience 31, 607–617 (2011).

21. Esfahani, M. J. et al. Closed-loop auditory stimulation of sleep slow oscillations: Basic principles and best practices. Neuroscience & Biobehavioral Reviews 153, 105379 (2023).

22. Alipour, M., Drongelen, W. van, Malerba, P., Voss, J. & Satzer, D. Predictability of Sleep Slow Oscillation Emergence and Spatial Extent from Pre-Onset Neural Dynamics. bioRxiv 2026–01 (2026).

23. Li, G. et al. Optimal referencing for stereo-electroencephalographic (SEEG) recordings. NeuroImage 183, 327–335 (2018).

24. Berry, R. B. et al. AASM scoring manual updates for 2017 (version 2.4). Journal of clinical sleep medicine 13, 665–666 (2017).

25. Ren, G. et al. Association between interictal high-frequency oscillations and slow wave in refractory focal epilepsy with good surgical outcome. Frontiers in Human Neuroscience 14, 335 (2020).

26. Alipour, M., Seok, S., Mednick, S. C. & Malerba, P. A classification-based generative approach to selective targeting of global slow oscillations during sleep. Frontiers in Human Neuroscience 18, 1342975 (2024).

27. Dang-Vu, T. T. et al. Spontaneous neural activity during human slow wave sleep. Proceedings of the National Academy of Sciences 105, 15160–15165 (2008).

28. Brunner, D. et al. Muscle artifacts in the sleep EEG: Automated detection and effect on all-night EEG power spectra. Journal of sleep research 5, 155–164 (1996).

29. Wang, C. et al. An attempt to identify reproducible high-density EEG markers of PTSD during sleep. Sleep 43, zsz207 (2020).

30. Daume, J., Gruber, T., Engel, A. K. & Friese, U. Phase-amplitude coupling and long-range phase synchronization reveal frontotemporal interactions during visual working memory. Journal of Neuroscience 37, 313–322 (2017).

31. Tort, A. B., Komorowski, R., Eichenbaum, H. & Kopell, N. Measuring phase-amplitude coupling between neuronal oscillations of different frequencies. Journal of neurophysiology 104, 1195–1210 (2010).

32. Berens, P. CircStat: a MATLAB toolbox for circular statistics. Journal of statistical software 31, 1–21 (2009).

33. Maris, E. & Oostenveld, R. Nonparametric statistical testing of EEG-and MEG-data. Journal of neuroscience methods 164, 177–190 (2007).

34. Malerba, P., Whitehurst, L. N., Simons, S. B. & Mednick, S. C. Spatio-temporal structure of sleep slow oscillations on the electrode manifold and its relation to spindles. Sleep 42, zsy197 (2019).

35. Sanchez-Vives, M. V. Origin and dynamics of cortical slow oscillations. Current Opinion in Physiology 15, 217–223 (2020).

36. Mak-McCully, R. A. et al. Coordination of cortical and thalamic activity during non-REM sleep in humans. Nature communications 8, 15499 (2017).

37. Kumar, V. J., Scheffler, K. & Grodd, W. The structural connectivity mapping of the intralaminar thalamic nuclei. Scientific Reports 13, 11938 (2023).

38. Willis Jr, W. D., Zhang, X., Honda, C. N. & Giesler Jr, G. J. A critical review of the role of the proposed VMpo nucleus in pain. The journal of pain 3, 79–94 (2002).

39. Van der Werf, Y. D., Witter, M. P. & Groenewegen, H. J. The intralaminar and midline nuclei of the thalamus. Anatomical and functional evidence for participation in processes of arousal and awareness. Brain research reviews 39, 107–140 (2002).

