## Supplementary Materials for "Thalamic Nuclei Differentially Coordinate Propagation of Cortical Slow Oscillations"

**Tables**

**Supplementary Table S1. Recording coverage by thalamic nucleus and patient.** For each nucleus, the number of patients, recording nights, and total contacts (NREM, W = 0.5 s coupling window) are shown. Per-patient columns show the number of recording contacts contributed by each patient to each nucleus; dashes indicate no recording in that patient. Nuclei with only one recording contact (ANT, VA) are included for completeness but were excluded from inferential analyses. Abbreviations: ANT, anterior nucleus; CL, central lateral; CM, centromedian; MD, mediodorsal; Pf, parafascicular; PuA, pulvinar anterior; PuL, pulvinar lateral; PuM, pulvinar medial; VA, ventral anterior; VL, ventral lateral; VPL, ventral posterolateral.

| Nucleus | Patients | Nights | Contacts | Patient ID | | | | | |
|---|---|---|---|---|---|---|---|---|---|
| | | | | 001 | 003 | 013 | 023 | 028 | 029 |
| ANT | 1 | 1 | 1 | — | — | — | — | 1 | — |
| CL | 1 | 6 | 6 | — | — | — | — | — | 6 |
| CM | 2 | 8 | 16 | — | 8 | — | 8 | — | — |
| MD | 1 | 6 | 6 | — | — | — | — | — | 6 |
| Pf | 1 | 4 | 4 | — | — | — | 4 | — | — |
| PuA | 1 | 4 | 4 | — | 4 | — | — | — | — |
| PuL | 5 | 18 | 21 | 6 | 4 | 6 | 4 | 1 | — |
| PuM | 5 | 18 | 78 | 24 | 16 | 18 | 16 | 4 | — |
| VA | 1 | 1 | 1 | — | — | — | — | 1 | — |
| VL | 1 | 4 | 8 | — | — | — | 8 | — | — |
| VPL | 3 | 14 | 32 | — | 4 | — | 4 | — | 24 |

**Supplementary Table S2. Temporal co-occurrence between thalamic nucleus SOs and PFC SO troughs across sleep stages.** Group-level peri-event histogram results (±1 s window, 50 ms bins) for all 11 thalamic nuclei plus pooled thalamus (All nuclei pooled) across N2 and N3 sleep stages separately. Contact count (N) = number of recording contacts per nucleus and stage. Peak Time (ms) = time of maximum co-occurrence probability relative to the PFC SO trough (time = 0; negative = thalamic SO precedes PFC trough; positive = thalamic SO follows PFC trough). Significant Bins = number of contiguous significant bins out of 40 total (cluster-based permutation test, 1,000 permutations, α = 0.05, two-tailed; cluster p reported). Blue shading indicates nuclei with statistically significant temporal coupling (cluster $p < 0.05$). ns = non-significant. Grey shading indicates nuclei with N = 1 (ANT, VA), for which group-level inference is not supported.

| Nucleus | Stage | N | Peak Time (ms) | Significant Bins (/ 40) | Cluster p |
|---|---|---|---|---|---|
| Pooled all nuclei | N2 | 24 | 150 | 40 / 40 | < 0.001 |
| Pooled all nuclei | N3 | 16 | 50 | 40 / 40 | < 0.001 |
| PuM | N2 | 78 | 150 | 40 / 40 | < 0.001 |
| PuM | N3 | 56 | 50 | 37 / 40 | < 0.001 |
| PuL | N2 | 21 | -100 | 40 / 40 | < 0.001 |
| PuL | N3 | 16 | 50 | 40 / 40 | < 0.001 |
| PuA | N2 | 4 | 150 | 0 / 40 | ns |
| PuA | N3 | 4 | 100 | 0 / 40 | ns |
| CM | N2 | 16 | -350 | 40 / 40 | < 0.001 |
| CM | N3 | 10 | -450 | 40 / 40 | < 0.001 |
| MD | N2 | 6 | -450 | 27 / 40 | 0.035 |
| MD | N3 | 2 | -350 | 0 / 40 | ns |
| VPL | N2 | 32 | -400 | 40 / 40 | < 0.001 |
| VPL | N3 | 8 | -300 | 40 / 40 | 0.008 |
| VL | N2 | 8 | -250 | 40 / 40 | 0.008 |
| VL | N3 | 8 | -450 | 40 / 40 | 0.011 |
| CL | N2 | 6 | -350 | 38 / 40 | 0.034 |
| CL | N3 | 2 | 450 | 0 / 40 | ns |
| ANT | N2 | 1 | 600 | 0 / 40 | ns |
| ANT | N3 | 1 | -500 | 0 / 40 | ns |
| VA | N2 | 1 | -50 | 0 / 40 | ns |
| VA | N3 | 1 | 50 | 0 / 40 | ns |
| Pf | N2 | 4 | -100 | 0 / 40 | ns |
| Pf | N3 | 4 | 50 | 0 / 40 | ns |

**Supplementary Table S3. Phase coupling between thalamic nucleus SOs and PFC SO troughs across sleep stages.** Group-level Rayleigh test results for all 11 thalamic nuclei plus pooled thalamus (All nuclei pooled) across N2 and N3 sleep stages separately, assessed within a ±0.5 s coupling window centered on each PFC SO trough. Contact count (N) = number of valid recording contacts per nucleus and stage. Group MVL = group mean vector length (0–1; higher values reflect stronger cross-unit phase consistency). Preferred Phase (°) = preferred phase of thalamic SO troughs relative to the PFC SO cycle (0° = PFC SO up-state peak [upper hemisphere]; ±180° = down-state trough [lower hemisphere]; negative values = ascending post-trough phase; positive values = pre-trough descending phase). Rayleigh Z = group-level Rayleigh statistic for circular non-uniformity. FDR correction was applied across nuclei within each sleep stage. Blue shading indicates nuclei with significant phase coupling (Rayleigh $p < 0.05$; FDR-corrected). Grey shading indicates nuclei with N = 1 (ANT, VA), for which MVL = 1.000 and Rayleigh Z = 1.000 are mathematical properties of single-observation circular statistics and should not be interpreted as evidence of phase coupling.

| Nucleus | Stage | N | Group MVL | Preferred Phase (°) | Rayleigh Z | p-value |
|---|---|---|---|---|---|---|
| Pooled all nuclei | N2 | 24 | 0.288 | +176.8° | 1.99 | 0.137 |
| Pooled all nuclei | N3 | 16 | 0.295 | -142.2° | 1.92 | 0.147 |
| PuM | N2 | 78 | 0.129 | -157.5° | 1.29 | 0.275 |
| PuM | N3 | 64 | 0.292 | -131.2° | 5.45 | 0.004 |
| PuL | N2 | 21 | 0.089 | -165.4° | 0.16 | 0.851 |
| PuL | N3 | 18 | 0.267 | +168.8° | 1.28 | 0.282 |
| PuA | N2 | 4 | 0.516 | -39.3° | 1.07 | 0.369 |
| PuA | N3 | 1 | 1.000 | -10.3° | 1.00 | 0.512 |
| CM | N2 | 16 | 0.275 | -152.7° | 1.21 | 0.303 |

| Nucleus | Stage | N | Group MVL | Preferred Phase (°) | Rayleigh Z | p-value |
|---|---|---|---|---|---|---|
| CM | N3 | 10 | 0.574 | -136.2° | 3.29 | 0.033 |
| MD | N2 | 6 | 0.695 | +11.9° | 2.90 | 0.048 |
| MD | N3 | 6 | 0.146 | -135.6° | 0.13 | 0.889 |
| VPL | N2 | 32 | 0.214 | -28.1° | 1.46 | 0.233 |
| VPL | N3 | 8 | 0.111 | +154.3° | 0.36 | 0.704 |
| VL | N2 | 8 | 0.386 | -167.4° | 1.19 | 0.313 |
| VL | N3 | 8 | 0.448 | -78.3° | 1.61 | 0.205 |
| CL | N2 | 6 | 0.285 | +23.3° | 0.49 | 0.635 |
| CL | N3 | 6 | 0.339 | +175.3° | 0.69 | 0.523 |
| ANT | N2 | 1 | 1.000 | +174.3° | 1.00 | 0.512 |
| ANT | N3 | 1 | 1.000 | -159.4° | 1.00 | 0.512 |
| VA | N2 | 1 | 1.000 | +8.8° | 1.00 | 0.512 |
| VA | N3 | 1 | 1.000 | +14.4° | 1.00 | 0.512 |
| Pf | N2 | 4 | 0.474 | -111.8° | 0.90 | 0.435 |
| Pf | N3 | 4 | 0.287 | -120.5° | 0.33 | 0.744 |

**Supplementary Table S4. Thalamocortical slow oscillation temporal and phase coupling during NREM sleep: Frontal slow oscillations.** Temporal and phase coupling between thalamic nucleus SOs and PFC SO troughs during pooled NREM sleep (N2+N3) for Frontal SOs. All column definitions as in Table 1. Blue shading indicates statistically significant results (cluster-based permutation test for temporal coupling; FDR-corrected Rayleigh test for phase coupling). Frontal SOs are associated with significant phase locking in pooled thalamus (All; Z = 3.545, p = 0.027, preferred phase = −156°) and PuM (Z = 5.174, p = 0.005, preferred phase = −170°), the strongest single phase coupling result across both SO types in the NREM dataset. The ascending post-trough preferred phases in both pooled thalamus and PuM indicate that thalamic SOs tend to co-occur as the cortex transitions from down-state back toward the up-state during Frontal SO events. Grey shading indicates nuclei with N = 1 (ANT, VA), for which MVL = 1.000 and Rayleigh Z = 1.000 are mathematical properties of single-observation circular statistics and should not be interpreted as evidence of phase coupling.

| Nucleus | Contact count | Temporal coupling | | Phase coupling | | |
|---|---|---|---|---|---|---|
| | | Max rate (%) | Max rate bin (ms) | Group MVL | Preferred Phase (°) | Rayleigh Z |
| Pooled all nuclei | 24 | 9.6315 | 150 | 0.3843 | -156.35 | 3.545 |
| PuM | 78 | 1.1752 | 150 | 0.2576 | -169.82 | 5.174 |
| PuL | 21 | 1.1406 | 150 | 0.251 | -177.59 | 1.323 |
| PuA | 4 | 1.9774 | 150 | 0.5212 | -52.73 | 1.086 |
| CM | 16 | 1.4205 | 150 | 0.2987 | -162.79 | 1.427 |
| MD | 6 | 1.745 | -100 | 0.4657 | 30.65 | 1.301 |
| VPL | 32 | 1.863 | 150 | 0.3258 | -130.49 | 0.849 |
| VL | 8 | 2.1829 | -700 | 0.567 | -134.62 | 2.572 |
| CL | 6 | 1.9786 | -300 | 0.4413 | 66.05 | 1.168 |
| ANT | 1 | 3.9691 | -350 | 1 | -144.74 | 1 |
| VA | 1 | 4.9063 | 50 | 1 | 24.19 | 1 |
| Pf | 4 | 1.8352 | 350 | 0.5131 | -158.66 | 1.053 |

**Supplementary Table S5. Thalamocortical slow oscillation temporal and phase coupling during NREM sleep: Global slow oscillations.** Temporal and phase coupling between thalamic nucleus SOs and PFC SO troughs during pooled NREM sleep (N2+N3) for Global SOs. All column definitions as in Table 1. Blue shading indicates statistically significant results (cluster-based permutation test for temporal coupling; FDR-corrected Rayleigh test for phase coupling). Grey shading indicates nuclei with N = 1 (ANT, VA), for which MVL = 1.000 and Rayleigh Z = 1.000 are mathematical properties of single-observation circular statistics and should not be interpreted as evidence of phase coupling. Notably, CM is the only nucleus showing significant phase coupling with Global SOs (Z = 4.136, FDR p = 0.013, preferred phase = +175°; near the down-state trough).

| Nucleus | Contact count | Temporal coupling | | Phase coupling | | |
|---|---|---|---|---|---|---|
| | | Max rate (%) | Max rate bin (ms) | Group MVL | Preferred Phase (°) | Rayleigh Z |
| Pooled all nuclei | 24 | 9.5163 | 50 | 0.1916 | -141.05 | 0.881 |
| PuM | 78 | 1.0879 | 150 | 0.1285 | -151.52 | 1.288 |
| PuL | 21 | 1.2266 | ~ (-100 to +50) | 0.1376 | -161.77 | 0.397 |
| PuA | 4 | 1.0389 | 250 | 0.4041 | -6.48 | 0.653 |
| CM | 16 | 1.3975 | -300 | 0.5084 | 175.02 | 4.136 |
| MD | 6 | 2.358 | -300 | 0.6019 | 5.51 | 2.174 |
| VPL | 32 | 1.3747 | -250 | 0.2695 | 178.2 | 0.581 |
| VL | 8 | 2.6934 | -450 | 0.2971 | -133.86 | 0.706 |
| CL | 6 | 2.9497 | 550 | 0.4718 | 48.38 | 1.336 |
| ANT | 1 | 4.3968 | -550 | 1 | -163.26 | 1 |
| VA | 1 | 11.1612 | 50 | 1 | 10.56 | 1 |
| Pf | 4 | 2.0078 | 50 | 0.6511 | -161.46 | 1.696 |

**Supplementary Table S6.** Phase coupling between thalamic nucleus SOs and PFC SO troughs for Global SOs across sleep stages. Group-level Rayleigh test results for Global SOs across all 11 thalamic nuclei plus pooled thalamus, for N2 and N3 sleep stages separately, assessed within a ±0.5 s coupling window centered on each PFC SO trough. Contact count (N) = number of recording contacts per nucleus and stage. FDR correction was applied across nuclei within each sleep stage. Blue shading indicates nuclei with significant phase coupling (Rayleigh $p < 0.05$; FDR-corrected). Grey shading indicates nuclei with N = 1 (ANT, VA), for which MVL = 1.000 and Rayleigh Z = 1.000 are mathematical properties of single-observation circular statistics and should not be interpreted as evidence of phase coupling.

| Nucleus | Stage | N | Grp MVL | Preferred Phase (°) | Rayleigh Z | p-value |
|---|---|---|---|---|---|---|
| Pooled all nuclei | N2 | 24 | 0.047 | -136.4° | 0.054 | 0.949 |
| Pooled all nuclei | N3 | 22 | 0.264 | -115.1° | 1.532 | 0.218 |
| PuM | N2 | 78 | 0.040 | -156.6° | 0.126 | 0.882 |
| PuM | N3 | 64 | 0.256 | -143.3° | 4.186 | 0.015 |
| PuL | N2 | 21 | 0.139 | -103.3° | 0.407 | 0.671 |
| PuL | N3 | 18 | 0.271 | +141.9° | 1.319 | 0.271 |
| PuA | N2 | 4 | 0.391 | -17.4° | 0.611 | 0.574 |
| CM | N2 | 16 | 0.437 | +179.8° | 3.051 | 0.045 |
| CM | N3 | 10 | 0.690 | -172.6° | 4.756 | 0.006 |
| MD | N2 | 6 | 0.611 | +9.1° | 2.243 | 0.103 |
| MD | N3 | 5 | 0.255 | -71.5° | 0.326 | 0.742 |
| VPL | N2 | 32 | 0.279 | -172.5° | 0.623 | 0.551 |
| VPL | N3 | 8 | 0.397 | +179.2° | 0.631 | 0.563 |
| VL | N2 | 8 | 0.197 | -156.4° | 0.312 | 0.744 |
| VL | N3 | 8 | 0.156 | -109.6° | 0.196 | 0.831 |
| CL | N2 | 6 | 0.347 | +24.3° | 0.723 | 0.505 |
| CL | N3 | 6 | 0.448 | +130.7° | 1.206 | 0.313 |
| ANT | N2 | 1 | 1.000 | +141.9° | 1.000 | 0.512 |
| ANT | N3 | 1 | 1.000 | -156.7° | 1.000 | 0.512 |
| VA | N2 | 1 | 1.000 | +7.8° | 1.000 | 0.512 |

| Nucleus | Stage | N | Grp MVL | Preferred Phase (°) | Rayleigh Z | p-value |
|---|---|---|---|---|---|---|
| VA | N3 | 1 | 1.000 | +13° | 1.000 | 0.512 |
| Pf | N2 | 4 | 0.759 | -109.9° | 2.303 | 0.095 |
| Pf | N3 | 4 | 0.064 | +179° | 0.016 | 0.986 |

**Supplementary Table S7. Phase coupling between thalamic nucleus SOs and PFC SO troughs for Frontal SOs across sleep stages.** Group-level Rayleigh test results for Frontal SOs across all 11 thalamic nuclei plus pooled thalamus, for N2 and N3 sleep stages separately, assessed within a ±0.5 s coupling window centered on each PFC SO trough. FDR correction was applied across nuclei within each sleep stage. Blue shading indicates nuclei with significant phase coupling (Rayleigh $p < 0.05$; FDR-corrected). Grey shading indicates nuclei with N = 1 (ANT, VA), for which MVL = 1.000 and Rayleigh Z = 1.000 are mathematical properties of single-observation circular statistics and should not be interpreted as evidence of phase coupling.

| Nucleus | Stage | N | Grp MVL | Preferred Phase (°) | Rayleigh Z | p-value |
|---|---|---|---|---|---|---|
| Pooled all nuclei | N2 | 24 | 0.320 | +173.7° | 2.461 | 0.084 |
| Pooled all nuclei | N3 | 22 | 0.579 | -158.4° | 7.375 | 0.000 |
| PuM | N2 | 78 | 0.188 | +156.8° | 2.770 | 0.062 |
| PuM | N3 | 64 | 0.389 | -162.6° | 9.541 | 0.000 |
| PuL | N2 | 21 | 0.269 | -176.9° | 1.518 | 0.221 |
| PuL | N3 | 18 | 0.487 | -166.3° | 4.264 | 0.012 |
| PuA | N2 | 4 | 0.521 | -55.5° | 1.086 | 0.362 |
| PuA | N3 | 1 | 1.000 | -23.6° | 1.000 | 0.512 |
| CM | N2 | 16 | 0.283 | -170.5° | 1.280 | 0.282 |

| Nucleus | Stage | N | Grp MVL | Preferred Phase (°) | Rayleigh Z | p-value |
|---|---|---|---|---|---|---|
| CM | N3 | 9 | 0.385 | -129.5° | 1.337 | 0.270 |
| MD | N2 | 6 | 0.679 | +16.7° | 2.767 | 0.056 |
| MD | N3 | 6 | 0.226 | +125.9° | 0.307 | 0.752 |
| VPL | N2 | 32 | 0.191 | -147.6° | 0.291 | 0.759 |
| VPL | N3 | 8 | 0.460 | +171.7° | 1.058 | 0.367 |
| VL | N2 | 8 | 0.521 | -145° | 2.175 | 0.112 |
| VL | N3 | 8 | 0.436 | -138.4° | 1.519 | 0.225 |
| CL | N2 | 6 | 0.512 | +64.2° | 1.574 | 0.214 |
| CL | N3 | 6 | 0.479 | +177.9° | 1.377 | 0.263 |
| ANT | N2 | 1 | 1.000 | -179.1° | 1.000 | 0.512 |
| ANT | N3 | 1 | 1.000 | -143.6° | 1.000 | 0.512 |
| VA | N2 | 1 | 1.000 | +20.7° | 1.000 | 0.512 |
| VA | N3 | 1 | 1.000 | +25° | 1.000 | 0.512 |
| Pf | N2 | 4 | 0.038 | -112.6° | 0.006 | 0.995 |
| Pf | N3 | 4 | 0.502 | +171.9° | 1.010 | 0.390 |

**Figures**

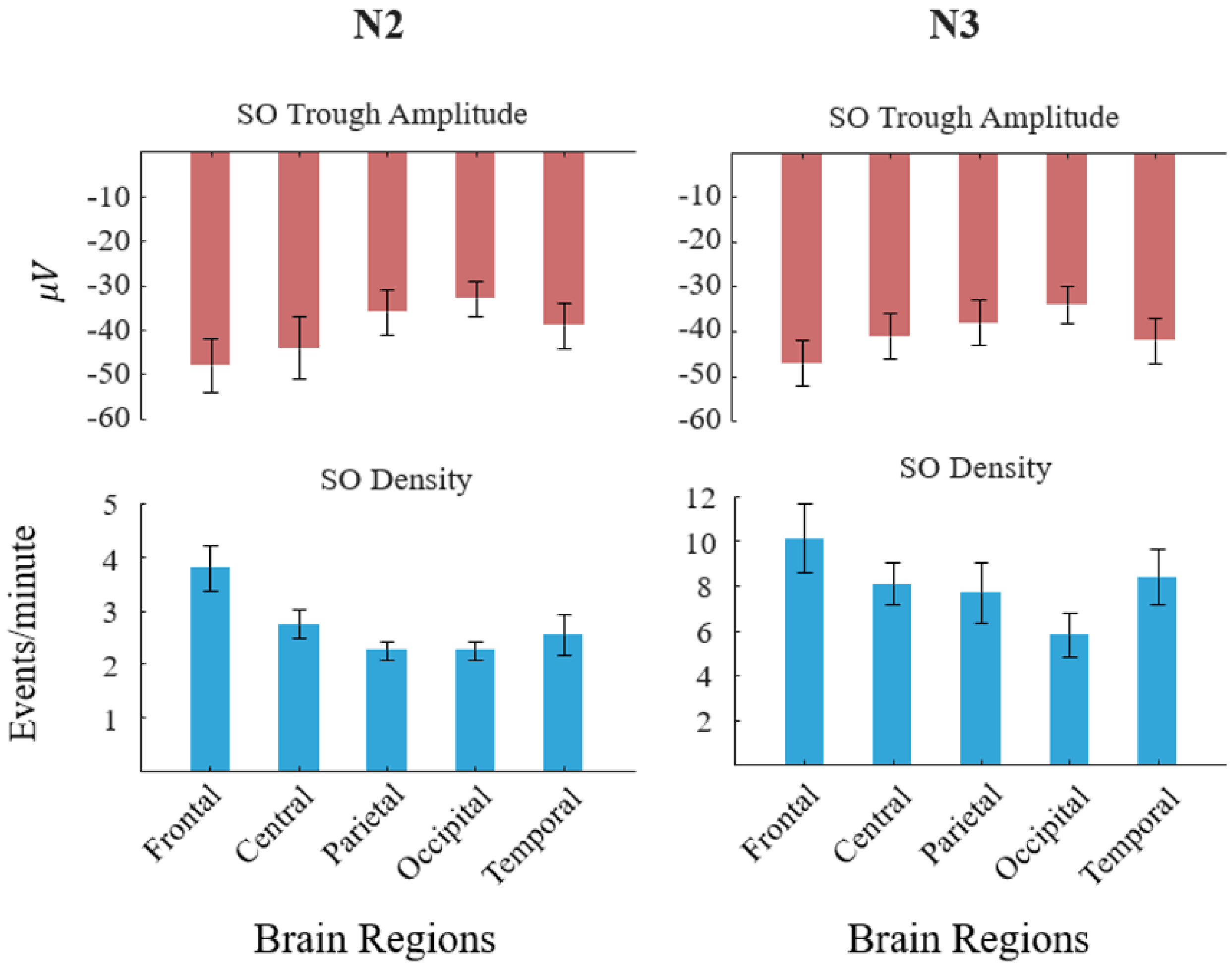


**Supplementary Figure S1. Scalp SO spatial characteristics separately for N2 and N3 sleep.** Mean (±SEM) SO trough amplitude (μV, upper panels) and SO rate (events/min, lower panels) across five scalp electrode regions (Frontal, Central, Parietal, Occipital, Temporal), shown separately for N2 (left column) and N3 (right column) sleep. Both stages show a clear frontal predominance in SO amplitude and density, consistent with the anterior cortical origin of canonical slow oscillations. N3 exhibits markedly higher SO density than N2, particularly at frontal electrodes, reflecting the deeper cortical synchrony of stage N3. The anteroposterior gradient in trough amplitude is preserved in both sleep stages.
